# USP9X promotes the degradation of trapped translation factors on collided ribosomes

**DOI:** 10.64898/2026.04.23.720364

**Authors:** Wojciech Teodorowicz, Lukas-Adrian Gurzeler, Oliver Mühlemann

## Abstract

Various ribosome-associated quality control pathways safeguard translation fidelity by detecting and resolving aberrant translation events. Recently, a pathway that senses stalled ribosomes with an occluded A-site has been described. The small molecules NVS1.1 and Ternatin-4 induce ribosome stalling by trapping eRF1 and eEF1A1, respectively, within the ribosomal A-site, thereby promoting their ubiquitination and proteasomal degradation. Additionally to the previously identified factors GCN1, RNF14, and RNF25, we identify here the deubiquitinase USP9X as a regulator of this pathway. In the absence of USP9X deubiquitinase activity, eRF1 is still ubiquitinated but fails to undergo efficient degradation. However, compared to wildtype cells, K6-linked ubiquitin chains accumulate on eRF1 in cells lacking USP9X activity, suggesting that USP9X trims K6-linked ubiquitin chains during a late step of substrate processing. Furthermore, clearance of trapped eRF1 engages a feedback mechanism involving 4EHP and the integrated stress response (ISR) to suppress translation, whereas impaired degradation prevents translational shutdown.

## Introduction

Translation of mRNA is a tightly regulated process that balances efficiency with fidelity.^1^ To ensure translational accuracy and maintain cellular protein homeostasis, cells have evolved surveillance mechanisms that monitor and resolve errors during translation.^2–4^ One such mechanism is ribosome-associated quality control (RQC), a conserved pathway that detects and eliminates incomplete nascent polypeptides resulting from ribosomal stalling on aberrant mRNAs.^5^ Ribosomal stalling can occur due to various factors, including improperly processed mRNA, defects in aminoacylated tRNAs, mutated or impaired translation factors, translation inhibitors, and environmental stresses such as oxidation or UV exposure.^6,7^ Stalled ribosomes often lead to collisions with trailing ribosomes.^8,9^ Ribosome collisions are a key signal for RQC activation^10^ initiating a cascade of regulatory events, including ubiquitination of ribosomal proteins,^11^ which facilitates the recruitment of specific RQC factors to resolve the stalled complex.^12–16^

Previous studies have identified a specialized branch of the RQC pathway that is activated by small molecules such as Ternatin-4^17^ and NVS1.1,^18^ which trap eukaryotic release factor 1 (eRF1) and elongation factor 1A1 (eEF1A1), respectively, in the ribosomal A-site. These trapping events induce ubiquitination of the ribosomal protein eS31 (RPS27a) by the E3 ligase RNF25, followed by RNF14-mediated ubiquitination and degradation of the trapped eRF1^18^ or eEF1A1^17^. The ribosomal collision sensor GCN1 plays a critical role in this process by bridging collided ribosomes and promoting RNF14 activity.^17,18^ Importantly, the GCN1–RNF14–RNF25 signaling axis is not restricted to drug-induced stalling; it is also engaged under physiological conditions, such as during formaldehyde-induced RNA-protein crosslinking, which triggers ribosome collisions and RNF14-mediated ubiquitination of the crosslinked proteins, targeting them for subsequent proteasomal degradation.^19,20^

The ubiquitin-proteasome system (UPS) and translation are closely interconnected processes.^21^ The interplay between the UPS and translation ensures that protein synthesis is precisely regulated and that protein quality is maintained.^22^ Deubiquitinating enzymes (DUBs), which reverse ubiquitination, are essential regulators of the UPS and play significant roles in maintaining translation fidelity and RQC.^23–27^ While approximately 100 DUBs have been identified in humans,^28^ the physiological roles and specific substrates remain poorly understood, with only a subset well-characterized to date.^29^ Given the broad role of DUBs in protein quality control mechanisms, their involvement in the occluded-A site RQC pathway remains unexplored.

Here, we identify the DUB USP9X as a regulator of the occluded A-site RQC. We show that USP9X is required for clearance of A-site-trapped eEF1A1 and eRF1, induced by Ternatin-4 and NVS1.1, respectively. In the absence of USP9X, eRF1 remains ubiquitinated upon NVS1.1 treatment and accumulates K6-linked ubiquitin chains, indicating that USP9X shapes ubiquitin chain architecture. Because K6-linked ubiquitin chains function as recognition signals for VCP, and VCP inhibition blocks eRF1 degradation, loss of USP9X is likely to impair efficient VCP-mediated extraction of ubiquitinated eRF1 and limit its proteasomal degradation. Failure to degrade eRF1 prevents engagement of a feedback mechanism that enforces translational shut-off via 4EHP (eIF4E2) and the integrated stress response (ISR).

## Results

### USP9X facilitates the degradation of eRF1 and eEF1A1 during occluded A-site RQC

In a previous study,^18^ two complementary approaches were employed to identify factors involved in the RQC response activated by NVS1.1: a genome-wide CRISPR knockout (KO) screen to identify genes that, when inactivated, confer resistance to high concentrations of NVS1.1, and a genome-wide siRNA screen to identify genes required for NVS1.1-mediated readthrough of a *Renilla* luciferase reporter gene. Both screens independently identified RNF14, RNF25, and GCN1 as top candidates involved in the response to NVS1.1.^18^ Notably, USP9X, which was not previously linked to the occluded A-site RQC pathway, also emerged as a significant hit in both screens (Figure 1A). This finding prompted us to investigate the role of USP9X in NVS1.1-induced eRF1 degradation.

**Figure 1.**
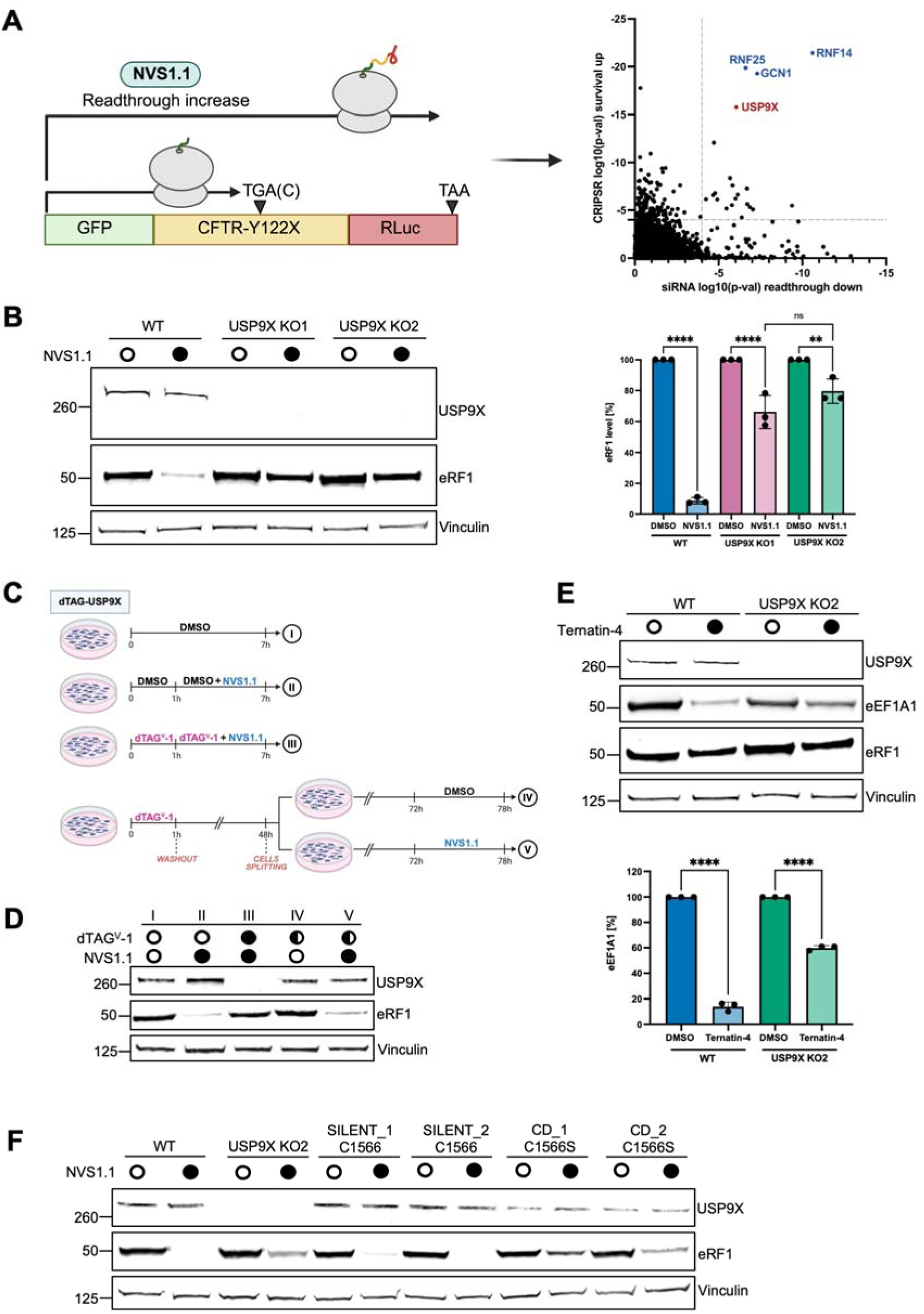
USP9X regulates the occluded A-site RQC pathway. **(A)** Schematic of the CFTR-Y122X-Rluc reporter used in genome-wide siRNA and CRISPR screens. Adapted from Gurzeler et al.^18^ **(B)** Immunoblot analysis and quantification of eRF1 levels in WT and USP9X KO cells treated with DMSO or NVS1.1 (25 μM, 6 h). eRF1 levels were quantified by densitometry and normalized to vinculin (mean ± SD, n = 3). Statistical analysis was performed using one-way ANOVA. **P < 0.01; ****P < 0.0001; ns, not significant. **(C)** Schematic of the dTAG-USP9X degradation and rescue strategy used in (D). **(D)** Immunoblot analysis of dTAG-USP9X cells treated with ± dTAGV-1 (500 nM, 1 h) and/or NVS1.1 (25 µM, 6 h). For the rescue experiment, cells were first treated with dTAGV-1 (500 nM, 1 h), washed with PBS, and incubated in dTAGV-1–free medium for 72 h (half circle), followed by treatment with DMSO or NVS1.1 at the same concentration and duration as above. **(E)** Immunoblot analysis and quantification of eEF1A1 levels in WT and USP9X KO cells treated with ternatin-4 (50 nM, 20 h). eEF1A1 levels were quantified by densitometry and normalized to vinculin (mean ± SD, n = 3). Statistical analysis was performed using ordinary one-way ANOVA. ****P < 0.0001. **(F)** Immunoblot analysis of WT, USP9X KO, and homozygous cell lines expressing either WT (silent) USP9X or a catalytically inactive (CD) USP9X mutant following treatment with NVS1.1 (25 μM, 6 h).

USP9X is a conserved DUB involved in multiple cellular processes, including translation,^30^ RQC,^31^ and apoptosis, ^32^ where it regulates distinct sets of proteins in a highly cell type-specific manner.^33^ To investigate its role in the pathway leading to NVS1.1-induced eRF1 degradation, we generated USP9X KO cells and treated them with NVS1.1. In agreement with previous findings,^18^ wild-type (WT) cells exhibited almost complete loss of eRF1 following treatment with NVS1.1 for 6 hours (Figure 1B). In contrast, USP9X KO cells showed impaired eRF1 degradation, though not a complete block (Figure 1B). The specificity of this phenotype was further validated by siRNA-mediated knockdown (KD), which recapitulated the reduced eRF1 degradation phenotype (Figure S1A). In an orthogonal approach, we treated WT cells with the selective USP9X inhibitor FT709, which has minimal activity against other DUBs.^31^ Consistent with the KO and KD experiments, FT709 treatment also partially inhibited eRF1 degradation following NVS1.1 treatment (Figure S1B), further supporting a specific role for USP9X in the occluded A-site RQC pathway.

As an additional validation, we established a reversible USP9X depletion system that enabled restoration of endogenous USP9X following transient depletion. An N-terminal dTAG degron was introduced into the endogenous USP9X *locus* by genome editing (Figure S1C).^34^ This approach allowed degradation of USP9X within 1 hour upon treatment with dTAG^V^-1 (Figure S1D), a heterobifunctional small molecule that binds the degron tag and recruits an E3 ubiquitin ligase, thereby inducing polyubiquitination of the tagged protein.^35^ We first examined whether acute depletion of USP9X phenocopied the KO. dTAG-USP9X cells were treated with dTAGV-1 for 1 hour to deplete USP9X or with DMSO as a control, followed by treatment with NVS1.1 for 6 hours (Figure 1C, conditions II and III). As expected, DMSO-treated cells exhibited almost complete degradation of eRF1 upon NVS1.1 treatment (Figure 1D, condition II), comparable to WT cells (Figure 1B), whereas acute depletion of USP9X markedly impaired eRF1 degradation (Figure 1D, condition III), recapitulating the USP9X KO phenotype.

We next asked whether restoration of USP9X could rescue NVS1.1-induced eRF1 degradation. USP9X was first depleted with dTAGV-1 for 1 hour, after which the compound was washed out, and cells were cultured in dTAGV-1-free medium for 72 hours to allow endogenous USP9X protein levels to recover before treatment with DMSO or NVS1.1 for 6 hours (Figure 1C, conditions IV and V). At this time point, USP9X expression was fully restored to levels comparable to untreated cells, and NVS1.1-induced eRF1 degradation was likewise restored (Figure 1D). Together, these findings demonstrate that efficient NVS1.1-mediated degradation of eRF1 requires USP9X.

Since Ternatin-4-induced degradation of eEF1A1 mirrors the mechanism of action of NVS1.1-induced eRF1 degradation,^18^ we investigated whether USP9X is also required for Ternatin-4-induced eEF1A1 degradation. As previously reported,^17^ Ternatin-4 treatment in WT cells led to a marked reduction in eEF1A1 protein levels. Under our experimental conditions, eEF1A1 levels decreased by approximately 85% following Ternatin-4 treatment (Figure 1E). However, in the USP9X KO cells, only a partial eEF1A1 degradation was observed to around 40% (Figure 1E). These findings suggest that USP9X functions more broadly in the degradation of A-site–trapped translation factors.

### USP9X’s catalytic activity is required for NVS1.1-mediated eRF1 degradation

USP9X contains an N-terminal ubiquitin-like (UBL) domain and a catalytic domain centered on the active-site cysteine C1566.^36,37^ The catalytic domain contains multiple ubiquitin-binding sites that enable recognition and processing of diverse polyubiquitin chains, including Lys6-, Lys11-, Lys48-, and Lys63-linked ubiquitin.^37,38^ To test whether the catalytic activity of USP9X is required for eRF1 degradation in response to NVS1.1, we generated two CRISPR KI cell lines: (i) a silent mutation in which the C1566 codon was altered without changing the encoded cysteine residue served as a control (Figure S1E), and (ii) a catalytically inactive mutant (C1566S) in which the cysteine codon was changed to a serine codon (Figure S1E). Upon NVS1.1 treatment, both WT and silent mutant cells exhibited a pronounced reduction in eRF1 levels (Figure 1F). By contrast, the catalytic-dead (CD) C1566S mutant and USP9X KO cells displayed only partial eRF1 degradation under the same conditions (Figure 1F), demonstrating that the enzymatic activity of USP9X is necessary for the efficient clearance of A-site trapped eRF1.

### Loss of USP9X leads to sustained eS31 ubiquitination during NVS1.1 exposure

A critical early event in the occluded A-site RQC pathway is the ubiquitination of the ribosomal protein eS31 at lysine 113 (K113) by the E3 ligase RNF25.^17,18^ This modification promotes the subsequent recruitment of RNF14 to stalled ribosomes via\ interaction with GCN1, which in turn ubiquitinates factors trapped in the A-site.^17,18^ Previous studies have shown that eS31 K113 ubiquitination increases ∼2-fold upon Ternatin-4 or NVS1.1 treatment,^17,18^ making this modification a useful marker for monitoring activation of the occluded A-site RQC pathway. We therefore asked whether the delayed eRF1 clearance observed in USP9X-deficient cells stems from altered eS31 ubiquitination.

Detection of eS31 ubiquitination in WT cells is challenging due to its rapid turnover and removal by the DUB USP16, which counteracts RNF25 activity.^39^ We therefore generated USP9X/USP16 double-knockout (dKO) cells to enhance detection of ubiquitinated eS31 following NVS1.1 treatment. In parallel, we also tested whether USP16 contributes to NVS1.1-induced eRF1 degradation. Given USP16’s antagonistic role toward RNF25, we hypothesized that it might enhance eRF1 ubiquitination and promote its turnover. However, NVS1.1 treatment for 1 or 6 hours led to comparable eRF1 degradation in WT and USP16 KO cells (Figure 2A), indicating that USP16 does not affect eRF1 stability. Moreover, eRF1 loss in the dKO closely mirrored that in USP9X KO cells (Figure 2A), arguing against any additive effect of USP16 deletion.

**Figure 2.**
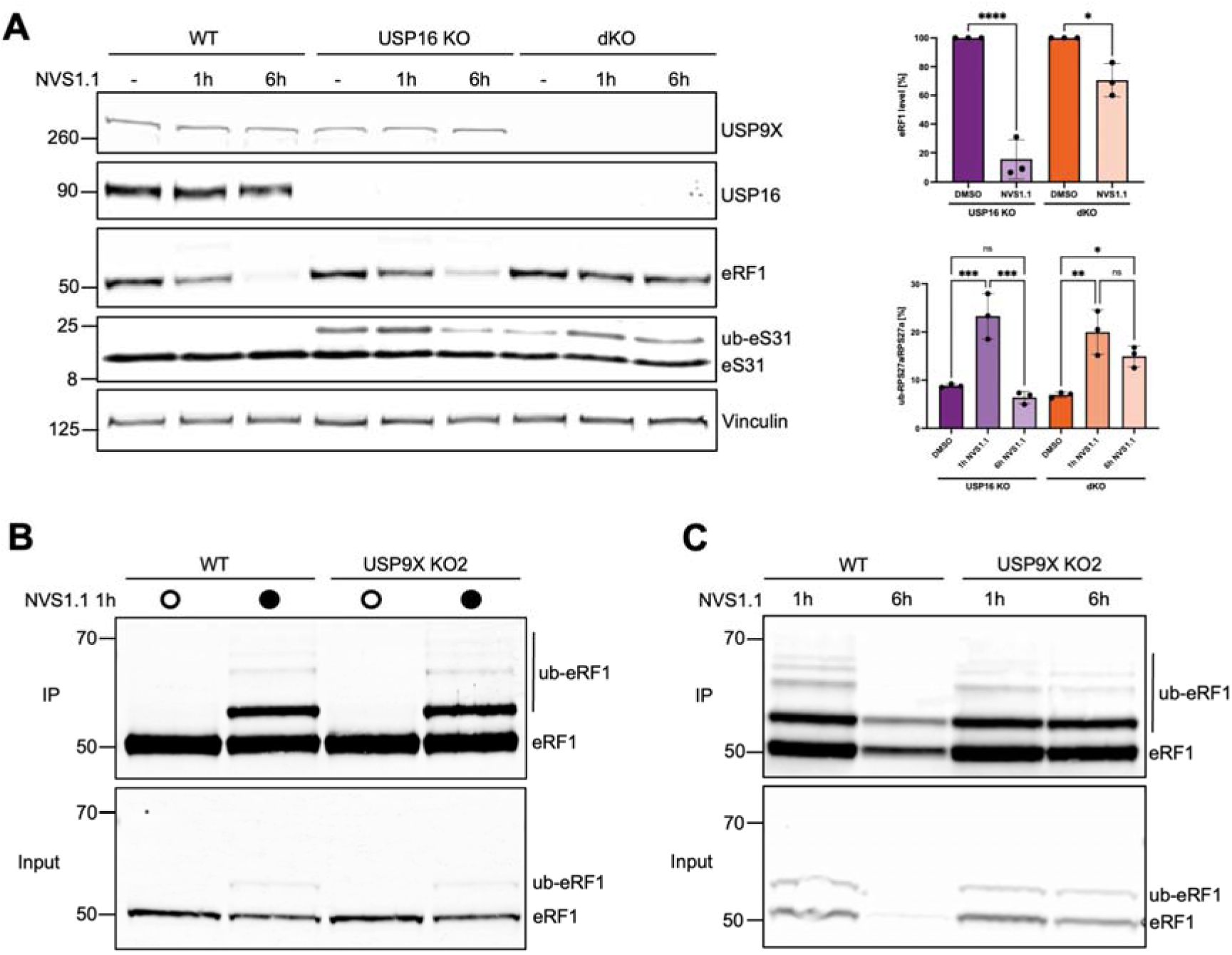
USP9X deficiency does not impair eRF1 ubiquitination or activation of the occluded A-site RQC pathway. **(A)** Immunoblot analysis and quantification of eRF1, eS31, and ubiquitinated eS31 in WT, USP16 KO, and USP9X/USP16 dKO cells treated with NVS1.1 (25 μM) for 1 or 6 h. eRF1 levels were quantified by densitometry and normalized to the loading control, and ubiquitinated eS31 was quantified relative to total eS31 (mean ± SD, n = 3). Statistical significance was determined using ordinary one-way ANOVA. ns, not significant; *P < 0.05; **P < 0.01; ***P < 0.001; ****P < 0.0001. **(B)** Immunoblot analysis of ubiquitinated eRF1 in WT and USP9X KO cells treated with DMSO or NVS1.1 (25 μM, 1 h). **(C)** Immunoblot analysis of ubiquitinated eRF1 in WT and USP9X KO cells treated with NVS1.1 (25 μM) for 1 or 6 h.

Next, eS31 ubiquitination was examined in cells treated with NVS1.1 or DMSO. In WT cells, no ubiquitinated eS31 was detected, regardless of NVS1.1 treatment (Figure 2A). However, USP16 KO cells showed two distinct bands corresponding to ubiquitinated and non-ubiquitinated eS31 (Figure 2A). Baseline ubiquitination accounted for ∼8% of total eS31, and increased to ∼20% after 1 hour of NVS1.1 treatment, consistent with prior reports^17^ (Figure 2A). Although loss of USP16 resulted in elevated basal eS31 ubiquitination compared to WT cells, this alone was insufficient to induce spontaneous eRF1 degradation in the absence of NVS1.1 treatment. This observation suggests that additional features, such as ribosome collisions and/or trapping of translation factors within the A-site, are required to trigger downstream degradation events.

After 6 hours of treatment with NVS1.1, ubiquitination returned to baseline, coinciding with reduced eRF1 levels (Figure 2A). In the dKO cells, 1-hour NVS1.1 treatment similarly induced a ∼2-fold increase in eS31 ubiquitination (Figure 2A). However, unlike in USP16 KO cells, this ubiquitination remained elevated after 6 hours, showing only a modest decline (∼1.5-fold above baseline) (Figure 2A).

Together, these data demonstrate that loss of USP9X does not impair the induction of eS31 ubiquitination but instead results in its prolonged persistence following NVS1.1 treatment, indicating that USP9X acts downstream of RNF25-mediated eS31 ubiquitination.

### USP9X depletion does not prevent eRF1 ubiquitination

Next, we investigated whether the KO of USP9X alters the NVS1.1-mediated ubiquitination of eRF1. We performed co-immunoprecipitation (co-IP) of eRF1 from WT and USP9X KO cells treated for 1 hour with NVS1.1 or DMSO. Surprisingly, no notable difference in eRF1 ubiquitination was observed between WT and USP9X KO cells (Figure 2B), indicating that loss of USP9X does not prevent eRF1 ubiquitination. To further examine whether USP9X affects the temporal dynamics of eRF1 ubiquitination, we repeated the co-IP experiments following both short (1 hour) and long (6 hours) NVS1.1 treatments. As expected, WT cells exhibited a strong reduction of eRF1 6 hours after NVS1.1 addition (Figure 2C). In contrast, eRF1 levels remained higher in USP9X KO cells, and the ubiquitination pattern was comparable between the 1-hour and 6-hour treatments (Figure 2C), suggesting that eRF1 remains ubiquitinated but is not efficiently degraded. Finally, to determine how long eRF1 ubiquitination persists in the absence of USP9X, we extended the NVS1.1 treatment duration to 12 and 24 hours. In WT cells, eRF1 levels were substantially reduced at these time points, and ubiquitinated eRF1 was hardly detectable (Figure S2). In contrast, USP9X KO cells retained ubiquitinated eRF1 even after 24 hours of treatment (Figure S2), consistent with persistent occluded A-site RQC activity.

### USP9X links eRF1 degradation to NVS1.1-induced translational repression

To this point, our data indicate that, in the absence of USP9X, NVS1.1 treatment still activates the occluded A-site RQC pathway, resulting in persistent eRF1 ubiquitination; however, its subsequent degradation is impaired. We first considered whether loss of USP9X affects proteasomal activity, thereby slowing eRF1 turnover. Proteasome activity was therefore measured in WT and USP9X KO cells, revealing no significant difference between the two (Figure S3A). We next evaluated an alternative explanation. Because the occluded A-site RQC pathway is a translation-dependent mechanism,^17,18^ and USP9X has been reported to regulate translation efficiency,^30^ we asked whether reduced translation is sufficient to impair eRF1 degradation. To address this, we partially inhibited global protein synthesis using the mTORC1/2 inhibitor Torin-1, which has been shown to suppress translation without completely blocking protein synthesis.^40^ Although Torin-1 substantially reduced translation, as confirmed by puromycin incorporation (Figure S3B), eRF1 degradation following NVS1.1 treatment remained unaffected (Figure S3B), indicating that a partial reduction in global translation is not sufficient to impair eRF1 degradation.

As NVS1.1 treatment induces both eRF1 degradation and translational repression, we next tested whether loss of USP9X also affects the translational response to NVS1.1. In WT cells, robust puromycin incorporation indicated active translation, whereas treatment with NVS1.1 for 6 hours completely abolished translation (Figure 3A). In contrast, both USP9X KO cell lines maintained active translation despite NVS1.1 treatment. A similar phenotype was observed following USP9X KD (Figure S3C), demonstrating that USP9X is required for NVS1.1-induced translational shutdown.

**Figure 3.**
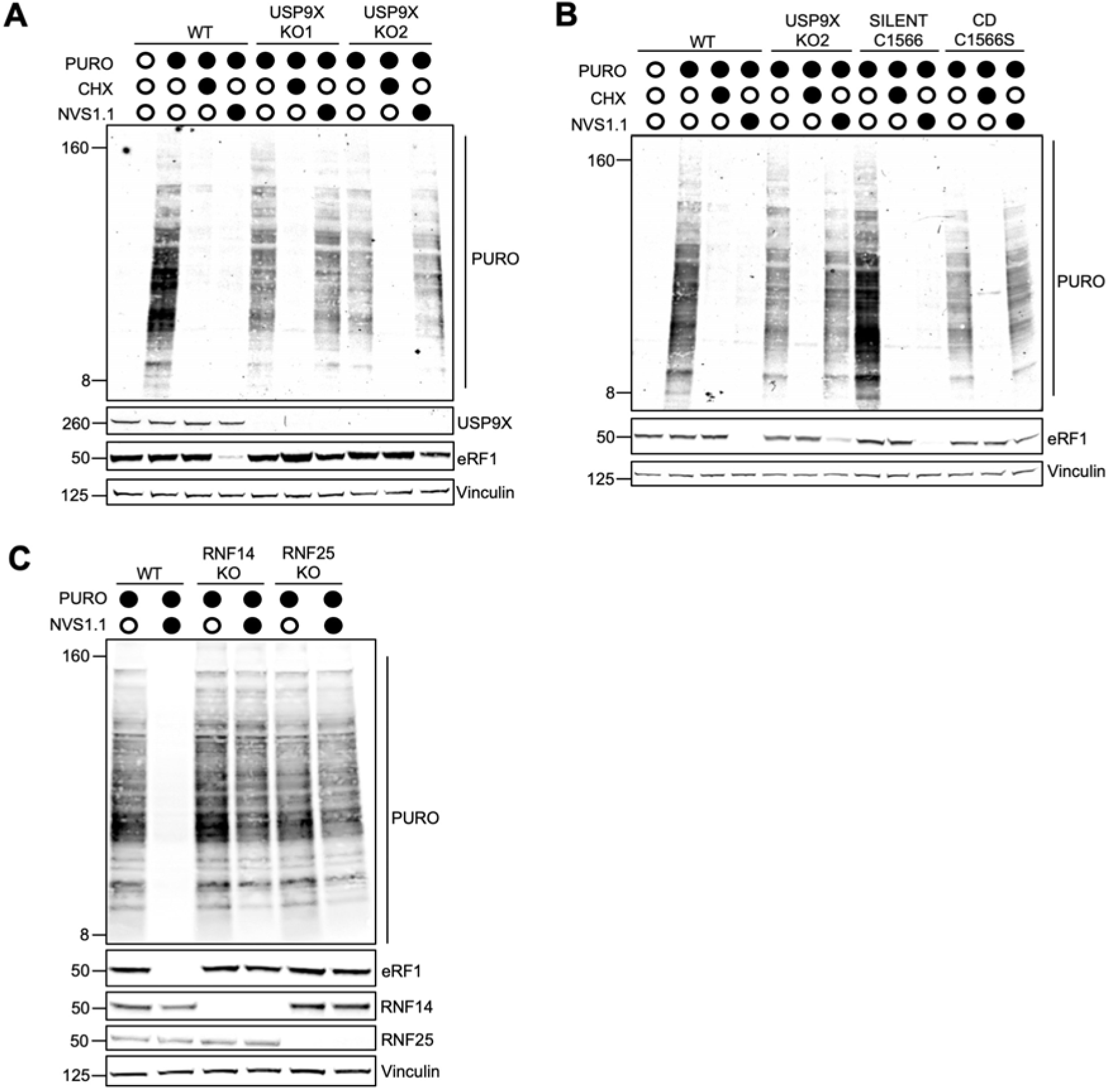
USP9X, RNF14, and RNF25 are required for NVS1.1-induced translational shutdown. **(A)** Puromycin incorporation assay measuring global protein synthesis in WT and USP9X KO cells. Cells were treated with DMSO or NVS1.1 (25 μM, 6 h), followed by puromycin (100 μg/mL, 15 min). Cycloheximide (100 μg/mL, 15 min) served as a translation inhibition control. Newly synthesized proteins were detected by immunoblotting with an anti-puromycin antibody. **(B)** Puromycin incorporation assay in WT, USP9X KO2, USP9X WT (silent), and catalytically inactive USP9X cell lines, performed as in (A). **(C)** Puromycin incorporation assay in WT, RNF14 KO, and RNF25 KO cells, performed as in (A).

To determine whether this function depends on the catalytic activity of USP9X, we performed puromycin incorporation assays in cells expressing either the silent or CD USP9X mutant. Whereas NVS1.1 efficiently suppressed translation in WT and silent mutant cells, CD cells phenocopied the USP9X KO, maintaining active translation despite NVS1.1 treatment (Figure 3B). These findings were confirmed using an orthogonal assay based on L-azidohomoalanine (L-AHA) incorporation. Consistent with the puromycin assay, NVS1.1 nearly abolished translation in WT and silent mutant cells but had a modest effect in CD cells (Figure S3D). Together, these results demonstrate that the deubiquitinase activity of USP9X is required for NVS1.1- induced translational repression.

Finally, we asked whether translational repression is uniquely dependent on USP9X or instead reflects a broader requirement for factors involved in eRF1 degradation. Puromycin incorporation assays performed in WT, RNF14 KO, and RNF25 KO cells showed that, similar to USP9X-deficient cells, both KO cell lines maintained active translation following NVS1.1 treatment (Figure 3C). These findings indicate that NVS1.1-induced translational repression depends on efficient clearance of trapped eRF1 rather than on USP9X alone.

### Loss of USP9X activity leads to accumulation of K6-linked ubiquitination on eRF1

The impaired eRF1 degradation observed following NVS1.1 treatment in cells expressing the CD USP9X mutant or lacking USP9X, together with the associated failure to induce translational shut-off, prompted us to identify potential USP9X substrates. USP9X has been implicated in the regulation of multiple interacting proteins by deubiquitinating and so stabilizing them, including eIF4A1,^30^ the E3 ligases ITCH,^41^ SMURF1,^42,43^ MARCH7,^42^ as well as two E3 ligases associated with RQC, Makorin2 and ZNF598.^31,44^ We first focused on recapitulating previously reported substrates, particularly eIF4A1 and ZNF598, as they are directly involved in translation and RQC, respectively. Under our experimental conditions, we did not observe a significant decrease in eIF4A1 levels in cells lacking USP9X (Figure S4A). However, we recapitulated the previously reported reduction in ZNF598 (Figure S4A,B). Consistent with this, loss of USP9X led to increased poly(A) readthrough (Figure S4C,D), suggesting impaired resolution of collided ribosomes via ZNF598. Moreover, combined depletion of ZNF598 and USP9X did not further exacerbate this effect (Figure S4D), indicating that these proteins function in the same pathway.

Nevertheless, because ZNF598 depletion does not affect NVS1.1-induced eRF1 degradation,^18^ it does not explain the involvement of USP9X in the occluded A-site RQC. Therefore, to systematically identify potential USP9X substrates, we performed diGly-based mass spectrometry in USP9X-active (USP9X_SILENT) and CD mutant cell lines following prolonged NVS1.1 treatment (6 hours), a condition in which eRF1 is efficiently degraded in the presence of catalytically active USP9X. This analysis revealed increased ubiquitination of previously reported USP9X substrates, including CEP131^45^ (at K110 and K320) and ITCH^41^ (at K405) (Figure 4A; Table S1 and Table S3). Ubiquitination sites on eIF4A1^30^ (K174 and K369) were also detected, but they were neither specifically enriched in the CD mutant nor did they reach statistical significance. Ubiquitination of various ribosomal proteins of the large subunit (LSU) was enriched in cells lacking USP9X activity, while ubiquitinated small subunit (SSU) proteins were detected both enriched and de-enriched (Figure 4A).

**Figure 4.**
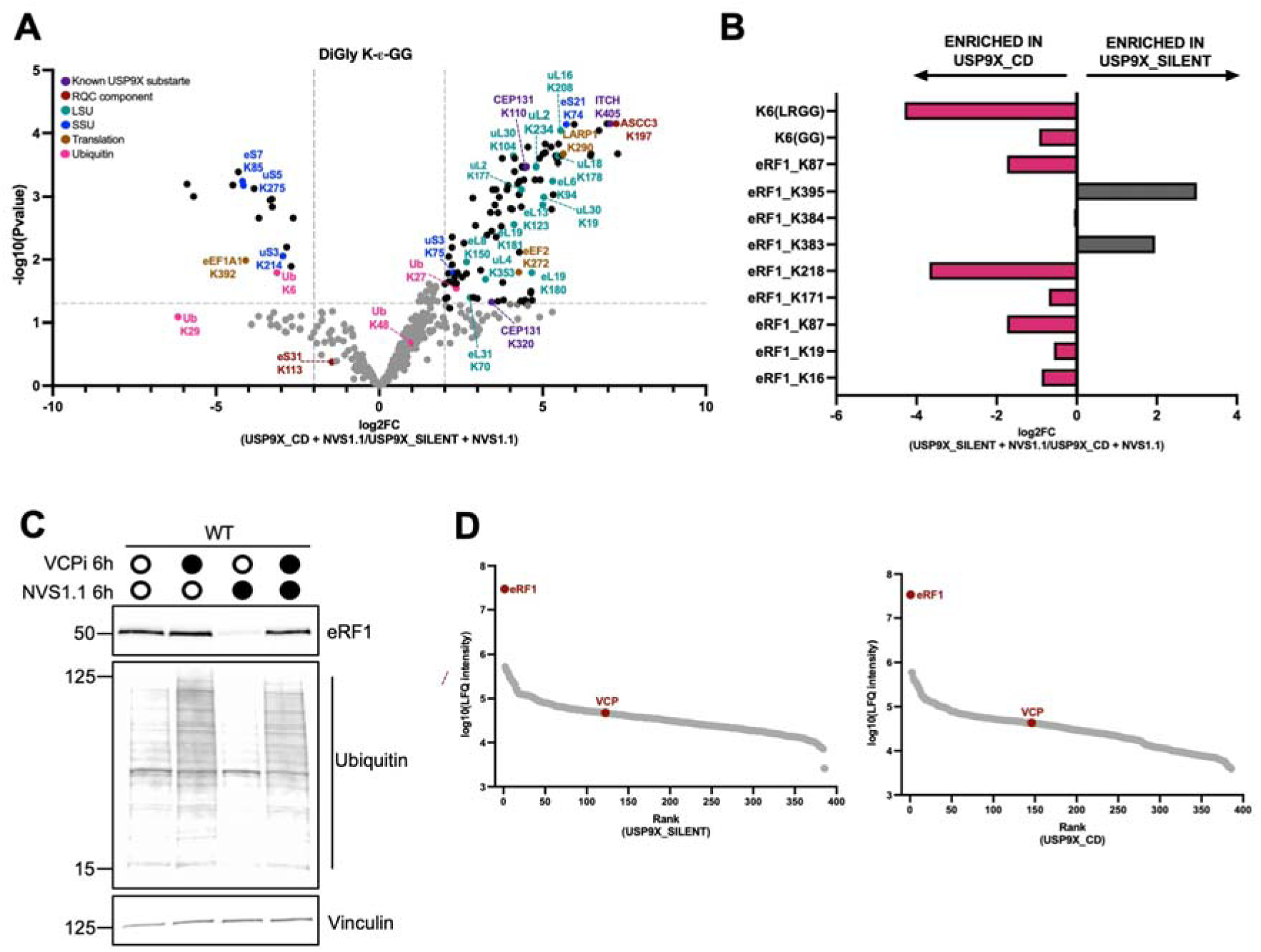
USP9X regulates K6-linked ubiquitination and VCP-dependent degradation of trapped eRF1. **(A)** DiGly proteomic analysis of USP9X WT (silent) and catalytically inactive USP9X cells treated with NVS1.1 (25 μM, 6 h). **(B)** DiGly proteomic analysis of eRF1 immunoprecipitates from USP9X WT (silent) and catalytically inactive USP9X cells treated with NVS1.1 (25 μM, 1 h). **(C)** Immunoblot analysis of cells treated with NVS1.1 (25 μM) in the presence or absence of a VCP inhibitor (10 μM, 6 h). **(D)** Rank plot of proteins identified by eRF1 immunoprecipitation from cells treated as in (B). Protein abundance is represented as the mean log10-transformed normalized label-free quantification (LFQ) intensity.

Analysis of ubiquitin linkage composition revealed an enrichment of K6-linked and an underrepresentation of K27-linked ubiquitination in NVS1.1-treated cells with active USP9X (USP9X_SILENT) compared to cells expressing catalytic-dead USP9X (Figure 4A). Because previous work demonstrated that SRI-41315 treatment,^17^ which also promotes eRF1 degradation,^46^ as well as Ternatin-4, both induce K6-linked ubiquitin chains,^17^ the altered K6-linked ubiquitin in USP9X_CD cells was of particular interest. Notably, K6-linked ubiquitination was shown to be abolished in RNF14 and RNF25 KO cells in response to Ternatin-4 treatment, supporting a role for these E3 ligases in assembling K6-linked chains.^17^ However, since eRF1 was not detected in the total lysate diGly dataset (Table S2), the observed increase in K6- linked ubiquitination could not be assigned to eRF1. We therefore next investigated whether K6-linked ubiquitination co-sedimented with collided ribosomes in polysome gradients and, ultimately, with eRF1 itself, and whether USP9X influences the ubiquitin chain architecture on eRF1.

First, we isolated nuclease-resistant collided ribosomes by polysome profiling coupled with micrococcal nuclease S7 digestion (Figure S5A). Proteomic analysis revealed significant enrichment of eRF1 and the collision marker EDF1 in heavy polysome fractions from NVS1.1-treated cells (Figure S5B; Table S4). Analysis of ubiquitin linkage composition further demonstrated a selective enrichment of K6- linked ubiquitination within these fractions compared with other ubiquitin linkage types (Figure S5C; Table S5).

Subsequently, ubiquitin linkage composition on eRF1 was analyzed by mass spectrometry following eRF1 immunoprecipitation from cells expressing either catalytically active or catalytic-dead USP9X after 1 hour of NVS1.1 treatment, when eRF1 ubiquitination patterns remain comparable between the two cell lines (Figure 2B). Mass spectrometry analysis identified multiple ubiquitination sites on eRF1 following NVS1.1 treatment, with K6-linked ubiquitination representing the predominant linkage type (Figure 4B; Table S6). K48-linked ubiquitination, which was 2-fold higher in cells with active USP9X (Table S6), was identified in only one sample from each condition and was therefore excluded from further analysis. Notably, K6- linked ubiquitination on eRF1 was enriched in cells expressing CD USP9X (Figure 4B), suggesting that USP9X contributes to trimming of K6-linked ubiquitin chains. Importantly, because K6-linked ubiquitination was detected in both catalytically active and inactive USP9X cells, RNF14-dependent assembly of K6-linked chains does not appear to require USP9X activity.

Together, these findings demonstrate that eRF1 is predominantly modified by K6- linked ubiquitin chains following NVS1.1 treatment and suggest that USP9X functions downstream of initial ubiquitination to trim K6-linked chains. Because loss of USP9X catalytic activity impairs efficient eRF1 degradation, USP9X-mediated deubiquitination may facilitate either the productive extraction of ubiquitinated eRF1 from the ribosomal A-site or its subsequent delivery to and degradation by the proteasome.

### eRF1 degradation is mediated by VCP

Ubiquitination commonly functions as a signal for proteasomal degradation. However, not all ubiquitinated proteins are readily accessible for proteasomal degradation. Proteins embedded within membranes or maintaining a folded conformation often require prior extraction or unfolding. This process is mediated by the AAA+ ATPase valosin-containing protein (VCP; also known as p97, Cdc48 in yeast, and TER94 in *Drosophila*), a conserved, chaperone-like enzyme that remodels ubiquitinated substrates before their downstream processing.^47^ K6-linked ubiquitin chains have been implicated in the recruitment of VCP during the clearance of RNA– protein crosslinks on stalled ribosomes in an RNF14-dependent manner.^19,20^ Although previous work established that eRF1 and eEF1A1 are degraded by the proteasome following NVS1.1^18^ and Ternatin-4 treatment,^17^ respectively, the requirement for VCP in this process had not been addressed. We investigated this possibility by treating WT cells with NVS1.1 in the presence or absence of a VCP inhibitor (VCPi). Co-treatment with VCPi blocked eRF1 degradation (Figure 4E), indicating that eRF1 turnover requires VCP activity.

Because inhibition of VCP or the proteasome could indirectly impair eRF1 degradation by preventing upstream eS31 ubiquitination, we examined eS31 ubiquitination in USP16 KO cells pretreated with either the VCP inhibitor (VCPi) or the proteasome inhibitor bortezomib (BTZ), followed by 1 hour of NVS1.1 treatment. In both conditions, eS31 ubiquitination remained readily detectable (Figure S6), indicating that neither inhibitor interferes with NVS1.1-induced eS31 ubiquitination. Thus, the effects of VCP and proteasome inhibition on eRF1 degradation occur downstream of eS31 ubiquitination.

Finally, we investigated VCP recruitment to ubiquitinated eRF1 by analyzing proteins co-immunoprecipitated with eRF1. VCP was detected in eRF1 immunoprecipitates from cells expressing either catalytically active or catalytically inactive USP9X (Figure 4D; Table S7), demonstrating that loss of USP9X catalytic activity does not impair VCP association with ubiquitinated eRF1.

Together, these findings indicate that VCP interacts with and is required for eRF1 degradation in NVS1.1-treated cells, whereas USP9X activity is dispensable for VCP- eRF1 interaction. This places USP9X function downstream of VCP-mediated extraction of ubiquitinated eRF1.

### NVS1.1 treatment engages ISR and RSR signalling with delayed kinetics

Although USP9X does not appear to directly mediate translational repression, the failure to suppress translation in USP9X-, RNF14-, and RNF25-deficient cells following NVS1.1 treatment prompted us to investigate the mechanism underlying translational shutdown. We first examined the kinetics of translational inhibition following NVS1.1 treatment and found that global translation was already strongly reduced after 2 hours (Figure 5A). To determine the effect of limiting eRF1 levels on ribosomes, we next monitored ribosome association with mRNAs by polysome profiling after 6 hours of NVS1.1 treatment, a condition in which translation is inhibited and eRF1 is largely degraded. Contrary to the expectation that impaired translation termination would increase ribosome occupancy on mRNAs and lead to polysome accumulation, we observed a marked loss of heavy polysomes accompanied by accumulation of 80S monosomes (Figure 5B). This phenotype is consistent with a recent study showing that acute degron-mediated depletion of eRF1 similarly reduces polysome abundance.^48^

**Figure 5.**
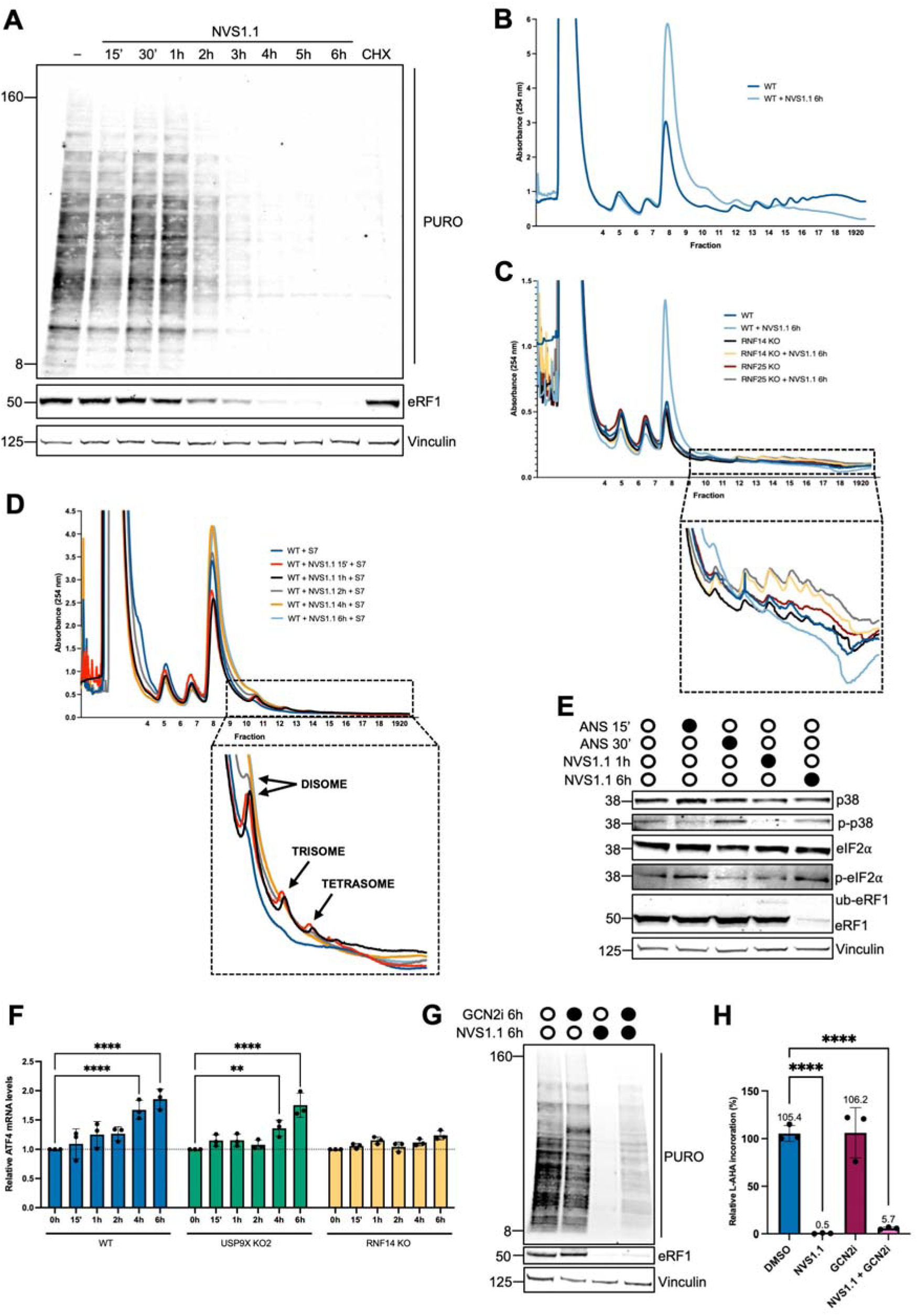
Ribosome collisions and translational shutdown precede ISR activation. **(A)** Time course of translational inhibition in WT cells treated with NVS1.1 (25 μM), measured by puromycin incorporation. **(B)** Polysome profiles of WT cells treated with DMSO or NVS1.1 (25 μM, 6 h). **(C)** Polysome profiles of WT, RNF14 KO, and RNF25 KO cells treated with DMSO or NVS1.1 (25 μM, 6 h). **(D)** Polysome profiles of WT cells treated with NVS1.1 (25 μM) for the indicated times (15 min–6 h). Cell lysates were treated with micrococcal nuclease (400 U, 40 min) before polysome profiling. **(E)** Immunoblot analysis of WT cells treated with anisomycin (1 mg/L, 15 min or 0.5 mg/L, 30 min) or NVS1.1 (25 μM, 1 or 6 h). **(F)** qPCR analysis of relative *ATF4* mRNA levels in WT, USP9X KO, and RNF14 KO cells treated with NVS1.1 (25 μM) for the indicated times. *ATF4* expression was normalized to *ACTB*. Data represent the mean ± SD of three independent biological replicates. Statistical significance was determined by two-way ANOVA. **, P < 0.01; ****, P < 0.0001. **(G)** Puromycin incorporation assay in WT cells treated with NVS1.1 (25 μM), a GCN2 inhibitor (1 μM), or both for 6 h. **(H)** Quantification of L-AHA incorporation measured as median fluorescence intensity (MFI), normalized to WT and expressed as relative L-AHA incorporation (%). Data represent mean ± SD (n = 3). Statistical significance was determined by ordinary one-way ANOVA. ****P < 0.0001.

If eRF1 depletion causes elongating ribosomes to accumulate within 3′ UTRs, the resulting reduction in available ribosomal subunits could limit translation initiation. We therefore examined polysome profiles in RNF14 KO and RNF25 KO cells, in which eRF1 ubiquitination and degradation are impaired. In contrast to WT cells, both Kos retained higher polysome levels following NVS1.1 treatment and, importantly, did not exhibit the marked accumulation of 80S monosomes observed in WT cells (Figure 5C). These findings indicate that sequestration of ribosomes within 3′ UTRs is not sufficient to explain the translational shutdown induced by NVS1.1.

To define the sequence of events leading to translational shutdown, we next examined the temporal dynamics of ribosome collisions following NVS1.1 treatment. Ribosome collisions were most abundant after 15 min and 1 hour, were markedly reduced by 2 hours, and were no longer readily detectable after 4 or 6 hours (Figure 5D), indicating that ribosome collisions represent an early and transient response to NVS1.1.

Acute degron-mediated depletion of eRF1 has been reported to activate the ISR,^48^ and extensive ribosome collisions activate both the ISR and the ribotoxic stress response (RSR).^15^ We therefore asked whether NVS1.1 activates these stress-response pathways, which in turn cause the translational shutdown. Activation of the ISR is mediated by GCN2-dependent phosphorylation of eIF2α, which reduces ternary complex formation and limits ribosome loading,^15,49–51^ whereas the RSR relies on ZAK -dependent activation of stress-activated protein kinases such as p38.^15^ While NVS1.1 treatment did not induce enhanced phosphorylation of either eIF2α or p38 after 1 hour, elevated levels of both p-eIF2α and p-p38 were observed after 6 hours of treatment (Figure 5E), indicating that activation of the ISR and RSR occurs only after the early phase of ribosome collisions.

WT, USP9X KO, and RNF14 KO cells differ in their ability to ubiquitinate and degrade eRF1 following NVS1.1 treatment, providing an opportunity to determine whether ISR activation is associated with eRF1 trapping, ubiquitination, or degradation. As a quantitative readout of ISR activation, we measured mRNA levels of the downstream effector *ATF4*. *ATF4* is a central transcriptional regulator induced during ISR activation and is also a well-characterized substrate of nonsense-mediated mRNA decay (NMD).^52^ As a positive control, cells were treated with the SMG1 inhibitor (SMG1i), which inhibits NMD.^53^ SMG1i treatment increased *ATF4* mRNA levels by approximately 2.6-fold (Figure S7). Following NVS1.1 treatment, *ATF4* mRNA levels were significantly increased after 4 hours in both WT and USP9X KO cells (Figure 5F). In contrast, no significant induction was observed in RNF14 KO cells regardless of NVS1.1 treatment (Figure 5F).

Finally, to determine whether activation of the ISR functionally contributes to the translational shutdown induced by NVS1.1, WT cells were treated with NVS1.1 in the presence or absence of a GCN2 inhibitor (GCN2i), which blocks phosphorylation of eIF2α. GCN2 inhibition minimally restored global translation after 6 hours of NVS1.1 treatment (Figure 5G and 5H). Nevertheless, eRF1 was still efficiently degraded (Figure 5G).

Collectively, these findings show that (i) ISR activation is not responsible for the early translational shutdown induced by NVS1.1, (ii) activation of the ISR occurs after ribosome collisions have largely resolved, indicating that these events do not temporally coincide, (iii) loss of USP9X activity does not prevent ISR activation in response to NVS1.1 despite impaired eRF1 degradation, and (iv) preventing both eRF1 ubiquitination and degradation, as observed in RNF14 KO cells, abolishes ISR activation.

### Translation initiation inhibition upon NVS1.1 treatment requires 4EHP

Canonical translation initiation in eukaryotes depends on eIF4E, which binds the 5′ cap of mRNAs and recruits additional initiation factors to assemble the translation initiation complex.^1^ A related cap-binding protein, 4EHP (also known as eIF4E2), shares structural similarity with eIF4E but instead acts as a translational repressor.^54^ 4EHP competes with eIF4E for cap binding and blocks access of the translation initiation complex to the mRNA.^55^ Given previous reports linking 4EHP activity to RQC,^8,56,57^ we hypothesized that the translation inhibition observed upon NVS1.1 treatment could be mediated by 4EHP. In contrast to WT, 4EHP KO cells retained high levels of translation following NVS1.1 treatment, indicating a loss of translational repression (Figure 6A). Interestingly, 4EHP KO cells also displayed only partial degradation of eRF1 upon NVS1.1 treatment (Figure 6A). To explore whether 4EHP and USP9X act in the same pathway, we depleted USP9X in 4EHP KO cells using siRNAs and assessed translation activity and eRF1 stability after NVS1.1 treatment. Combined depletion did not produce an additive effect (Figure 6A). Together, these results show that 4EHP is required for translational repression downstream of the occluded A-site RQC activation.

**Figure 6.**
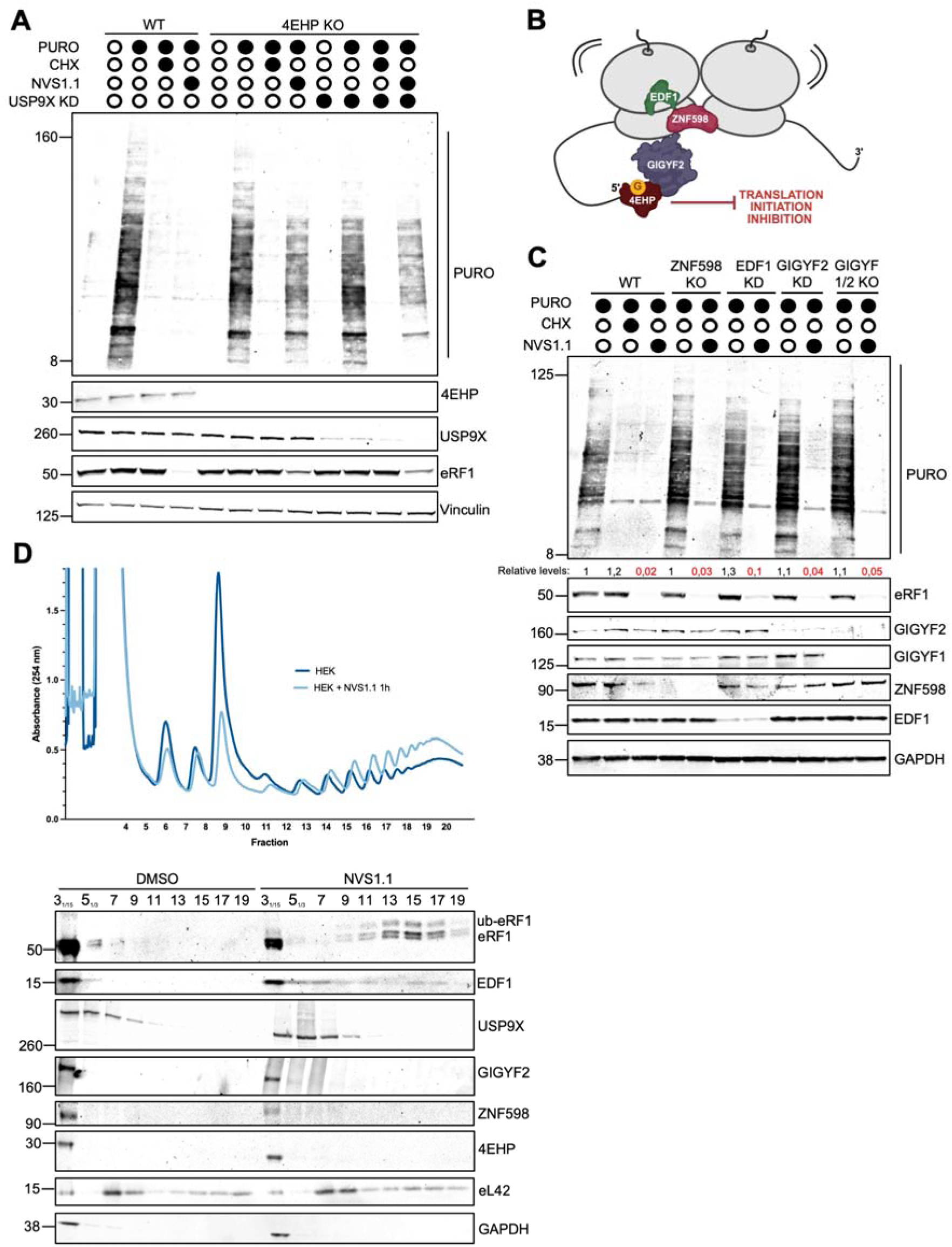
Translation inhibition downstream of the occluded A-Site RQC requires 4EHP. **(A)** Puromycin incorporation assay in WT, 4EHP KO, and 4EHP KO cells following USP9X KD, performed as in Figure 3A. **(B)** Schematic model of translational repression downstream of ribosome collision **(C)** Puromycin incorporation assay in WT, ZNF598 KO, EDF1 KD, GIGYF2 KD, and GIGYF2 KO cells. **(D)** Polysome profiles of cells treated with DMSO or NVS1.1 (25 μM, 1 h). Immunoblot analysis of proteins from odd-numbered fractions is shown below. Fractions 3 and 5 were diluted 1:15 and 1:3, respectively.

### Translation inhibition in the occluded A-site RQC is independent of GIGYF1/2, ZNF598, and EDF1

Unlike general translation repressors, 4EHP is recruited to specific transcripts by RNA-binding proteins, thereby exerting RNA-selective translational control.^55^ Notably, 4EHP forms a repressor complex with GIGYF2, a scaffold protein that stabilizes its interaction and helps recruit additional silencing factors.^55^ The 4EHP–GIGYF2 complex has been implicated in translation initiation inhibition of mRNAs with collided ribosomes, to which it is recruited through interaction with the zinc finger protein ZNF598,^55,56^ which in turn interacts with collided ribosomes and initiates protein and mRNA quality control pathways.^58,59^ This process is sensed by EDF1, which has been proposed to stabilize collided ribosomes and create a platform for recruitment of the ZNF598–GIGYF2–4EHP complex (Figure 6B).^8,57^ In addition to GIGYF2, GIGYF1, a paralog with overlapping function, has also been reported to interact with 4EHP and mediate translational repression in a 4EHP-dependent manner.^60^ To determine whether ZNF598, EDF1, or GIGYF1/2 contribute to the translational repression caused by NVS1.1, we evaluated global translation by puromycylation assays in ZNF598 KO, EDF1 KD, GIGYF2 KD, and GIGYF1/2 dKO cells. Despite efficient depletion of each factor, NVS1.1 treatment still caused translation shutdown in all cases (Figure 6C), indicating that none of these factors is essential for the NVS1.1-induced translation inhibition. Consistently, eRF1 was efficiently degraded upon NVS1.1 treatment in all conditions (Figure 6C), further supporting the conclusion that ZNF598, EDF1, and GIGYF1/2 are dispensable for both translational repression and eRF1 clearance.

Ribosome collisions have been proposed to act as a hub for recruiting the 4EHP– GIGYF2 complex, enabling repression of translation initiation in *cis* on specific transcripts.^57,61^ This model is supported by observations that low-dose emetine treatment, which induces ribosome collisions by stochastically stalling only a fraction of the elongating ribosomes, results in the accumulation of GIGYF2, ZNF598, and EDF1 in the fractions of polysome gradients that contain collided ribosomes.^61^ We next analyzed the distribution of 4EHP, GIGYF2, and ZNF598 across polysome gradients following 1 hour of NVS1.1 treatment (Figure 6D), a time point at which ribosome collisions are readily detectable. In agreement with previous observations^18^, eRF1 and EDF1 co-migrated with heavy polysome fractions corresponding to collided ribosomes upon NVS1.1 treatment (Figure 6D). In contrast, 4EHP, GIGYF2, and ZNF598 were detected exclusively in lighter, ribosome-free fractions of the sucrose gradient (Figure 6D). USP9X displayed a similar distribution and was not detectably altered by NVS1.1 treatment (Figure 6D). This may reflect either recruitment below the detection limit of our immunoblotting or a lack of stable association of these factors with collided ribosomes following NVS1.1-induced conditions.

## Discussion

RQC comprises multiple surveillance pathways that detect and resolve aberrant translation events. Among these, the occluded A-site response monitors ribosomes stalled due to persistent A-site occupancy. However, how this pathway interfaces with the ubiquitin–proteasome system and coordinates broader translational control remains largely uncharacterized. Here, we identify USP9X as an effector of the occluded A-site response. Loss of USP9X impairs complete eRF1 clearance following NVS1.1 treatment, attenuates Ternatin-4–induced degradation of eEF1A1, and prevents efficient translation arrest.

At first glance, the requirement for USP9X catalytic activity in the degradation of ubiquitinated eRF1 and eEF1A1 appears counterintuitive. However, USP9X is not the only DUB reported to facilitate efficient proteasomal degradation of ubiquitinated substrates.^62^ A similar role has been proposed for Ataxin-3 in endoplasmic reticulum-associated degradation (ERAD), where Ataxin-3 acts downstream of VCP-mediated retrotranslocation to trim polyubiquitin chains and maintain an optimal ubiquitin signal for proteasomal targeting.^63^ This remodeling step was proposed to coordinate substrate extraction with efficient proteasomal recognition.^63^ Additional support for such a mechanism came from studies of YOD1, another ERAD-associated DUB, whose dominant-negative mutant (YOD1 C160S) blocks substrate degradation.^64^ More recently, the yeast YOD1 ortholog Otu1 was shown to function directly with Cdc48, where substrate deubiquitination is coupled to Cdc48 activity.^65^ This process requires DUB-mediated trimming of polyubiquitinated substrates to enable their release from Cdc48 for subsequent degradation.^65^ Similarly, USP9X deficiency does not impair eRF1 ubiquitination *per se*, as evidenced by sustained eRF1 ubiquitination in USP9X-deficient cells upon NVS1.1 treatment, but instead alters ubiquitin chain architecture. This is reflected by the accumulation of K6-linked ubiquitin chains on eRF1 in cells expressing the catalytically inactive USP9X mutant. RNF14 was identified as the E3 ligase that specifically assembles K6-linked ubiquitin chains on eRF1 and eEF1A1 in response to SRI-41315 and Ternatin-4, respectively.^17^ In addition, RNF14 has been shown to catalyze K6 ubiquitination on crosslinked RNA-binding proteins.^20^ Previous studies demonstrated that K6-linked ubiquitin chains promote VCP-dependent processing of formaldehyde-induced RNA–protein crosslinks, enabling substrate extraction and proteasomal degradation.^19,20^ Consistent with this, we find that eRF1 degradation upon NVS1.1 treatment requires VCP activity, supporting a model in which K6-linked ubiquitination facilitates efficient clearance of trapped eRF1. Importantly, VCP association with eRF1 was detected in cells expressing both catalytically active and inactive USP9X, suggesting that substrate recognition by VCP is not impaired in the absence of USP9X activity, but downstream eRF1 processing and degradation are compromised. Together, these observations raise the possibility that, similarly to Ataxin-3 or YOD1/Otu1, USP9X functions downstream of initial ubiquitination to trim K6-linked ubiquitin chains and thereby promote productive extraction and proteasomal degradation of trapped eRF1.

Impaired clearance of eRF1 from the A-site has direct consequences for translational control. Persistent A-site occupancy prevents activation of feedback mechanisms that suppress translation initiation. Given that defective clearance of trapped factors directly impacts translational control, we examined how this response is coupled to global inhibition of protein synthesis. We find that translational repression in response to NVS1.1 depends on 4EHP, a cap-binding protein that competes with eIF4E to inhibit cap-dependent initiation.^56,66–68^ Interestingly, loss of 4EHP impairs efficient eRF1 degradation following NVS1.1 treatment, suggesting that translation shutdown facilitates clearance of trapped A-site factors. In the context of RQC, 4EHP has been reported to function via the GIGYF2–4EHP complex, recruited by adaptor proteins such as ZNF598 and EDF1.^56,61^ Yet, this pathway accounts for only a fraction of translational repression events,^66,67,69^ suggesting that 4EHP can also act independently of these adaptors. Consistent with this, GIGYF1/2, ZNF598, and EDF1 are dispensable for translational shutdown following NVS1.1 treatment. Together, our observations indicate that 4EHP mediates translational repression through an alternative mechanism. Because NVS1.1 targets ribosomes at stop codons present on virtually all mRNAs, the resulting inhibition is inherently global rather than transcript-specific. However, how 4EHP effectively competes with eIF4E1 to drive translational repression in response to NVS1.1, and whether additional cofactors contribute to this process, remain to be determined.

Our temporal analyses show that ISR and RSR activation occur only after prolonged NVS1.1 treatment, when collided ribosomes have largely been resolved, eRF1 has been degraded, and translation inhibition is maintained through a 4EHP-dependent mechanism. This temporal separation suggests that ISR and RSR activation are secondary consequences of sustained translational perturbation rather than immediate responses to ribosome collisions. It also raises the question of how cells transition from an early collision-induced quality-control response to later stress signaling. A mechanistic explanation may lie in the dynamic interactions of GCN1 with its binding partners. GCN1 is a conserved ribosome-associated scaffolding protein that binds the RWD domain of GCN2.^70,71^ In response to amino acid stress, GCN1 is required for GCN2 activation and subsequent induction of the ISR.^72,73^ In addition, GCN1 has been shown to associate with collided, ribosomes in both yeast^74^ and human cells,^15^ and to interact with the RWD domain of RNF14,^17^ providing an essential signaling input, most likely by recruiting and activating RNF14 near the occluded A-site of stalled ribosomes.^17,18^ Interestingly, recent studies showed that RNF25, which also contains an RWD domain, suppresses GCN2-dependent ISR activation through ubiquitination of eS31.^75–77^. However, how eS31 ubiquitination selectively promotes RNF14 function while limiting GCN2 activation remains unclear.^77^ One proposed mechanism is that ubiquitination of eS31 at K113, positioned near the RWD-binding region of GCN1, contributes to this selective signaling outcome. In this model, elevated eS31 ubiquitination may create a platform that preferentially stabilizes RNF14 association while restricting GCN2 engagement and downstream ISR signaling.^17^

This model may also help explain the differences between NVS1.1-induced translational repression and acute eRF1 depletion. A recent study linked eRF1 depletion to ISR-driven translational repression.^48^ In contrast to our observations, translational repression upon acute eRF1 depletion is independent of 4EHP.^48^ It is possible that during early stages of NVS1.1 treatment, when extensive ribosome collisions and eS31 ubiquitination are present, 4EHP-mediated translational repression represents the dominant response. As collided ribosomes are progressively resolved, and eS31 ubiquitination declines, ISR activation may emerge as a secondary response to reduced eRF1 levels. Accordingly, acute eRF1 depletion may resemble a later stage of the NVS1.1 response.

Collectively, we propose a model in which USP9X functions at a late step of the occluded A-site RQC pathway. Following RNF14-mediated ubiquitination and VCP recruitment, USP9X catalytic activity likely trims K6-linked ubiquitin chains to promote efficient processing and proteasomal degradation of trapped eRF1 or eEF1A1 (Figure 7). Whether USP9X acts by facilitating substrate release from VCP, analogous to deubiquitinases operating in ERAD, remains to be determined. Efficient clearance of trapped eRF1 or eEF1A1 is then coupled to 4EHP-dependent inhibition of translation initiation, thereby limiting further ribosome loading onto mRNAs containing occluded A-sites. Sustained translational repression subsequently engages the ISR, representing a secondary adaptive response rather than the primary driver of translational shutdown.

**Figure 7.**
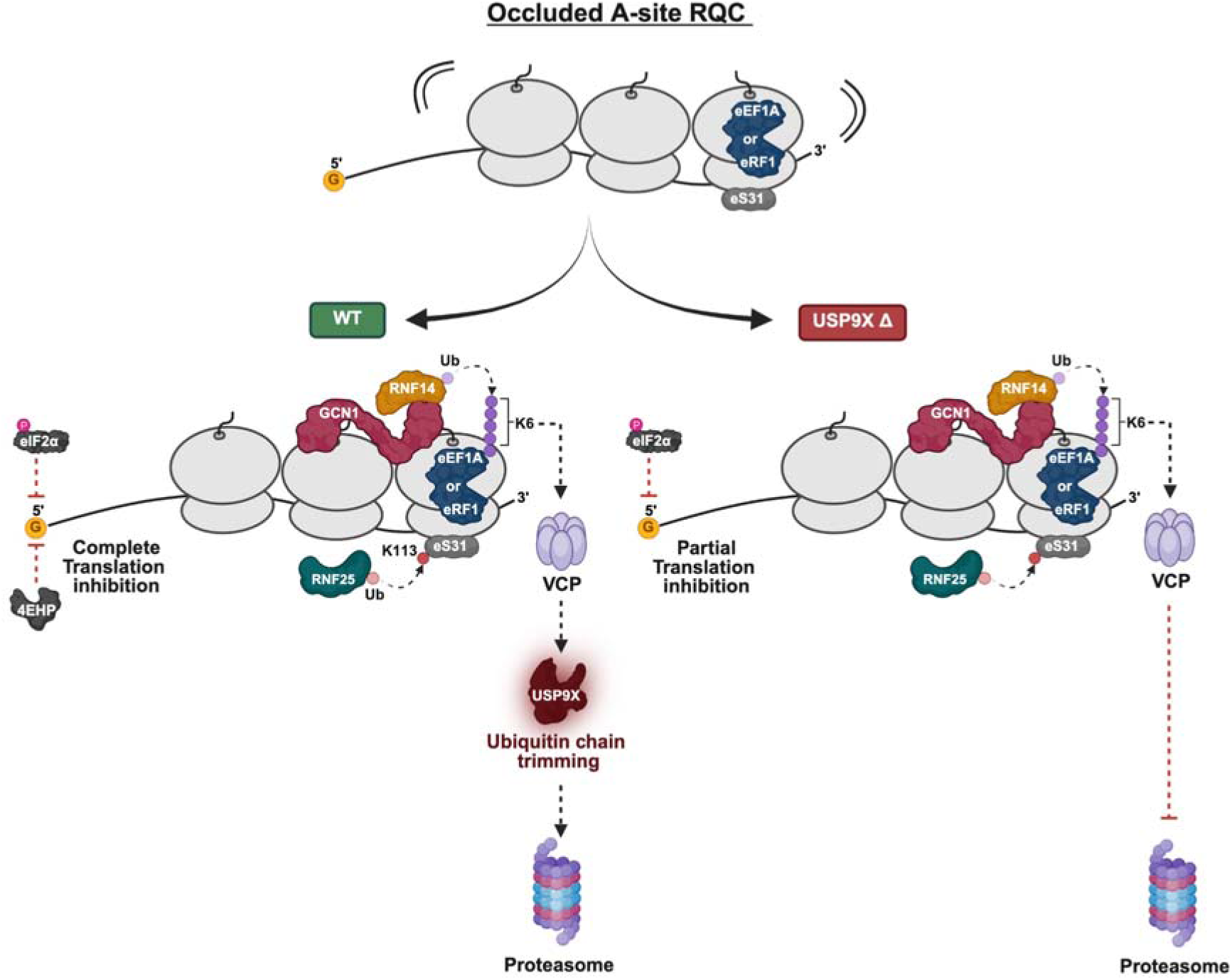
Proposed model of USP9X function in occluded A-site RQC.

### Limitations of the study

Our study examined the role of USP9X in the context of pharmacologically induced occluded A-site stress using NVS1.1 and Ternatin-4. Therefore, whether USP9X also contributes to the physiological clearance of endogenous substrates remains unclear. A reported physiological trigger of this pathway is the formaldehyde-induced stress response, which resolves RNA–protein crosslinks (RPCs).^19,20^ Given that RNF14, RNF25, GCN1, and K6-linked ubiquitination also function in RPC resolution, USP9X may likewise contribute to this process; however, this possibility remains to be experimentally addressed. We also note that the catalytically inactive USP9X protein is expressed at lower levels than wild-type or USP9X-silent variants. Nevertheless, the highly similar phenotypes observed in USP9X CD and USP9X KO cells argue that the defects primarily arise from loss of USP9X catalytic activity rather than reduced protein abundance. Finally, the mechanism linking the occluded A-site RQC to global translation repression remains incompletely understood. In particular, how 4EHP is recruited to mediate translation initiation inhibition, and why canonical 4EHP adaptors such as GIGYF2 are dispensable in this context, remain unresolved.

## Acknowledgments

We thank Jack Taunton for providing Ternatin-4, Jürgen Reinhardt (Novartis, Basel) for the generous gift of NVS1.1, and Ulrike Kutay for providing the eS31 antibody. We thank Sophie Braga Lagache, Natasha Buchs, Anne-Christine Uldry, and Manfred Heller from the Proteomics and Mass Spectrometry Core Facility at the University of Bern for excellent mass spectrometry support. We also thank Niels Gehring for providing the plasmids used to generate the dTAG knock-in, as well as Nicole Kleinschmidt and Karin Schranz for outstanding technical assistance. Figures 1A, 1C, 6B, and 7 were created using BioRender.

## Author contributions

O.M., W.T., and L.A.G. conceptualization. W.T. writing – original draft. W.T., O.M., and L.A.G. writing – review & editing. W.T. investigation. W.T. and O.M. data analysis & interpretation. W.T. visualization. O.M. funding acquisition. O.M. supervision.

## Declaration of generative AI and AI-assisted technologies in the writing process

During the preparation of this work, the authors used ChatGPT-5.2 to make the final manuscript more concise. After using this tool, the authors reviewed and edited the wording as needed and take full responsibility for the content of the publication.

## CONTACT FOR REAGENT AND RESOURCE SHARING

Further information and requests for reagents should be directed to the lead contact, Oliver Mühlemann.

## DATA AND CODE AVAILABILITY

- The MS raw data have been deposited in the PRIDE repository (http://www.ebi.ac.uk/pride) under the accession number: PXD079916
- This paper does not report original code.
- Any additional information required to reanalyze the data reported in this work paper is available from the lead contact upon request.

## SUPPLEMENTAL INFORMATION

Document S1. Figures S1–S7.

Table S1. Identification of diGly-modified peptides by mass spectrometry, related to Figure 4A.

Table S2. Total protein abundance analysis by mass spectrometry, related to Figure 4A.

Table S3. Relative abundance of diGly-modified peptides normalized to corresponding protein abundance, related to Figure 4A.

Table S4. Total protein mass spectrometry analysis of disome and heavy polysome fractions isolated from polysome profiles of cells treated with DMSO or NVS1.1 (1 h), related to Figures S5A and S5B.

Table S5. Identification of diGly-modified peptides in disome and heavy polysome fractions isolated from polysome profiles of cells treated with DMSO or NVS1.1 (1 h), related to Figures S5A and S5C.

Table S6. Identification of diGly-modified peptides from eRF1 immunoprecipitates by mass spectrometry, related to Figure 4B.

Table S7. Total protein mass spectrometry analysis of eRF1 immunoprecipitates, related to Figure 4D.

Table S8. Oligonucleotide sequences, related to STAR Methods.

## METHODS

### Key Resources Table

| REAGENT or RESOURCE | SOURCE | IDENTIFIER |
| --- | --- | --- |
| **Antibodies** |  |  |
| anti-eRF1, rabbit (immunoblot, 1:3'000) | Abcam | Cat# ab31799;<br>RRID:AB_732264 |
| anti-eRF1, mouse (IP) | Santa Cruz<br>Biotechnology | Cat# sc-365686;<br>RRID:AB_10843214 |
| anti-GAPDH, mouse (1:2'000) | Santa Cruz<br>Biotechnology | Cat# sc-47724;<br>RRID:AB_627678 |
| anti-vinculin, mouse (1:1'000) | Santa Cruz<br>Biotechnology | Cat# sc-73614;<br>RRID:AB_1131294 |
| anti-eEF1A1, rabbit (1:3'000) | Proteintech | Cat# 11402-1-AP;<br>RRID:AB_10232890 |
| anti-USP9X, mouse (1:1'000) | Santa Cruz<br>Biotechnology | Cat# sc-365353;<br>RRID:AB_10846088 |
| anti-eL42, mouse (1:1'000) | Santa Cruz<br>Biotechnology | Cat# sc-100831;<br>RRID:AB_2238666 |
| anti-ubiquitin (P4D1), mouse (1:1'000) | Santa Cruz<br>Biotechnology | Cat# sc-8017;<br>RRID:AB_628423 |
| anti-USP16, rabbit (1:1'000) | Bethyl Laboratories | Cat# A301-614A;<br>RRID:AB_1211385 |
| anti-eS31, rabbit (1:3'000) | Montellese et al.<sup>39</sup> | N/A |
| anti-puromycin clone 12D10, mouse (1:15'000) | Milipore | Cat# MABE343;<br>RRID:AB_2566826 |
| anti-p-eIF2γ (Ser51), rabbit (1:1'000) | Cell Signalling<br>Technology | Cat# 9721;<br>RRID:AB_330951 |
| anti-eIF2γ, mouse (1:1'000) | Cell Signalling<br>Technology | Cat# 2103;<br>RRID:AB_836874 |
| anti-p38 rabbit (1:1'000) | Cell Signalling<br>Technology | Cat# 9212;<br>RRID:AB_330713 |
| anti-p-p38 (Thr180/Tyr182) mouse (1:2'000) | Cell Signalling<br>Technology | Cat# 9216;<br>RRID:AB_331296 |
| anti-4EHP, rabbit (1:1'000) | Proteintech | Cat# 12227-1-AP;<br>RRID:AB_10642945 |
| anti-eIF4A, rabbit (1:1'000) | Cell Signalling<br>Technology | Cat# 2490;<br>RRID:AB_823487 |
| anti-GIGYF2, rabbit (1:10'000) | Proteintech | Cat# 24790-1-AP;<br>RRID:AB_2879727 |
| anti-GIGYF1, rabbit (1:1'000) | Proteintech | Cat# 31073-1-AP;<br>RRID:AB_3716590 |
| anti-ZNF598, rabbit (1:10'000) | Bethyl Laboratories | Cat# A305-108A;<br>RRID:AB_2631503 |
| anti-EDF1, rabbit (1:1'000) | Proteintech | Cat# 12419-1-AP;<br>RRID:AB_2230895 |
| anti-RNF14, rabbit (1:1'000) | Sigma-Aldrich | Cat# HPA008716;<br>RRID:AB_1856337 |
| anti-RNF25, rabbit (1:1'000) | Bethyl Laboratories | Cat# A303-844A;<br>RRID:AB_2620195 |
| IRDye® 800CW Donkey anti-Rabbit IgG (1:10'000) | LiCor | Cat# 926-32213 |
| IRDye® 680LT Donkey anti-Rabbit IgG (1:10'000) | LiCor | Cat# 926-68023 |
| IRDye® 680LT Donkey anti-Mouse IgG (1:10'000) | LiCor | Cat# 926-68022 |
| IRDye® 680LT Goat anti-Mouse IgG (1:10'000) | LiCor | Cat# 926-68050 |
| **Bacterial and virus strains** |  |  |
| Stellar competent cells | TakaRa | Cat# 636766 |
| Chemicals, peptides, and recombinant proteins |  |  |
| Cycloheximide | Focus Biomolecules | Cat# 10-117 |
| NVS1.1 | Gurzeler et al.<sup>18</sup> | N/A |
| Ternatin-4 | Oltion et al.<sup>17</sup> | N/A |
| Anisomycin | MedChemExpress | Cat# HY-18982 |
| Emetine | Cayman Chemical | Cat# TRC-E521530 |
| GCN2iB | MedChemExpress | Cat# HY-112654 |
| Puromycin | Sigma | Cat# P9620 |
| dTAG<sup>V</sup>-1 | MedChemExpress | Cat# HY-145514D |
| VCP/p97 inhibitor-1 | MedChemExpress | Cat# HY-139606 |
| SMG1 inhibitor (hSMG-1) | ProbeChem | Cat# PC-35788 |
| USP9X inhibitor (FT709) | Cayman Chemical | Cat# 39971 |
| Torin-1 | UBPBio | Cat# F6100 |
| Tet-free FBS | Gibco | Cat# A4736301 |
| Triton X-100 | Sigma-Aldrich | Cat# 93443 |
| TRIS buffer | Qiagen | Cat# 19086 |
| Trypsin-EDTA (0.25%) | Gibco | Cat# 25200056 |
| Trypsin | Promega | Cat# V5111 |
| PBS | Gibco | Cat# 10010001 |
| TWEEN® 20 | Sigma-Aldrich | Cat# P1379 |
| Phosphatase Inhibitor Cocktail I | TargetMol | Cat# C0002 |
| Phosphatase Inhibitor Cocktail II | TargetMol | Cat# C0003 |
| Protease Inhibitor Cocktail | TargetMol | Cat# 0001 |
| Ubiquitin Remnant (K-ε-GG) Immunoaffinity Beads | Cayman Chemical | Cat# 34699 |
| 4x NuPAGE LDS loading buffer | Thermo Fisher Scientific | Cat# NP0008 |
| Dynabeads™ Protein G | Invitrogen | Cat# 10004D |
| 1.5x LDS loading Bolt™ LDS Sample Buffer | Invitrogen | Cat# B0008 |
| RNase inhibitor | Vazyme | Cat# R301-3 |
| Micrococcal nuclease S7 | Thermo Fisher Scientific | Cat# EN0181 |
| Lipofectamine 2000 | Thermo Fisher Scientific | Cat# 11668019 |
| Lullaby | OZ Biosciences | Cat# LL71000 |
| TaKaRa Ex-Premier DNA Polymerase | Takara Bio | Cat# RR370A |
| EGTA | Thermo Fisher Scientific | Cat# 50-255-956 |
| DTT | Indofine Chemicals | Cat# MB1015 |
| DMEM (high glucose, no glutamine, no methionine, no cysteine) | Gibco | Cat# 21013024 |
| Opti-MEM reduced-serum medium | Gibco | Cat# 31985062 |
| L-Cysteine | Sigma-Aldrich | Cat# C7755 |
| GlutaMAX Supplement (100x) | Gibco | Cat# 35050-061 |
| Copper(II) sulfate (CuSO<sub>4</sub>) | Sigma-Aldrich | Cat# C1297 |
| Sodium L-ascorbate | Sigma-Aldrich | Cat# A7631 |
| Paraformaldehyde | Sigma-Aldrich | Cat# 158127 |
| L-Azidohomoalanine (AHA) | Vector Labs | Cat# CCT-1066 |
| Phosphate-Buffered Saline (PBS) (10x) | Thermo Fisher Scientific | Cat# J62036.K2 |
| XFD488 alkyne | AAT Bioquest | Cat# 1700 |
| Bovine Serum Albumin | Sigma-Aldrich | Cat# 05470 |
| Penicillin-Streptomycin (10,000 U/mL) | Invitrogen | Cat# 15140122 |
| N-ethylmaleimide (NEM) | MedChemExpress | Cat# HY-D0843 |
| FCS | Bioconcept | Cat# 2-01F16-I |
| HEPES | Invitrogen | Cat# 15630056 |
| DMEM | Invitrogen | Cat# 41966052 |
| CaCl<sub>2</sub> | Merck | Cat# 2382 |
| MgCl<sub>2</sub> | Fluka | Cat# 63063 |
| D-PBS | Invitrogen | Cat# 14190-144 |
| LysC | Promega | Cat# VA117A |
| Alt-R S.p Cas9 nuclease | IDT | Cat# 1081059 |
| Alt-R HDR Enhancer V2 | IDT | Cat# 10007921 |
| AffinityScript Multi-Temp Reverse Transcriptase | Agilent Technologies | Cat# 600109 |
| Brilliant III Ultra-Fast SYBR Green qPCR Master Mix | Agilent Technologies | Cat# 600883 |
| Critical commercial assays |  |  |
| In-Fusion cloning | Takara Bio | Cat # 638916 |
| Trans-Blot Turbo RTA Midi 0.2 mm PVDF Transfer Kit | Bio-Rad | Cat# 1704273 |
| Maxwell® RSC Genomic DNA Kit | Promega | Cat# AS1880 |
| Maxwell® RSC simplyRNA Cells Kit | Promega | Cat# AS1390 |
| Proteasome Assay kit | Cayman chemicals | Catt# 10008041 |
| TotalStain Q (NC) | Azure | Cat# AC2227 |
| Deposited data |  |  |
| Label-free mass spectrometry data | This study | PRIDE: PXD079916 |
| Experimental models: Cell lines |  |  |
| HEK293 FlpIn T-REx | Montellese et al.<sup>39</sup> | N/A |
| HEK293 FlpIn T-REx _USP16_KO | Montellese et al.<sup>39</sup> | N/A |
| HEK293 FlpIn T-REx _USP9X_KO | This study | N/A |
| HEK293 FlpIn T-REx _dKO (USP16 and USP9X) | This study | N/A |
| HEK293 FlpIn T-REx _USP9X_CD_C1566S | This study | N/A |
| HEK293 FlpIn T-REx _USP9X_silent_C1566C | This study | N/A |
| HEK293 FlpIn T-REx _ZNF598_KO | Garzia et al.<sup>58</sup> | N/A |
| HEK293 FlpIn T-REx _dKO (ZNF598 and USP9X) | This study | N/A |
| HEK293 FlpIn T-REx _dTAG-USP9X | This study | N/A |
| HEK293T | Peter et al.<sup>60</sup> | N/A |
| HEK293T_4EHP_KO | Peter et al.<sup>60</sup> | N/A |
| HEK293T_dKO_GIGYF1/2 | Peter et al.<sup>60</sup> | N/A |
| HEK293T_RNF14 KO | Gurzeler et al.<sup>18</sup> | N/A |
| HEK293T_RNF25 KO | Gurzeler et al.<sup>18</sup> | N/A |
| Oligonucleotides |  |  |
| Primer, siRNA, sgRNA sequences | This paper | See Table S8 |
| Recombinant DNA |  |  |
| USP9X_pSpCas9_BB-2A-Puro (p2318) | This study | N/A |
| GFP_P2A_K(AAA)20-P2A_RFP (p2437) | Addgene | Cat# 105688 |
| GFP_P2A_RFP (p2438) | Addgene | Cat# 105686 |
| pCRIS-PITChv2 | Addgene | Cat# 91796 |
| pX330-BbsI-PITCh | Addgene | Cat# 127875 |
| pCRIS-PITChv2_USP9X (p2581) | This study | N/A |
| pX330-BbsI-PITCh_USP9X (p2587) | This study | N/A |
| Software and algorithms |  |  |
| GraphPad | GraphPad Software | Software version 9.0 |
| MaxQuant | Max-Planck Institute of Biochemistry | Software version 1.3.0.5 |
| FragPipe | Nesvilab | Software version 1.8 |
| IonQuant | Nesvilab | Software version 1.8 |
| Biorender | Biorender | https://www.biorender.com |
| FlowJo | BD Biosciences | https://flowjo.com |
| SnapGene | SnapGene software | www.snapgene.com |
| Other |  |  |
| Trans-Blot Turbo Transfer System | Bio-Rad | Cat# 1704150<br>Cat# 1704158 |
| Imaging System | LiCor | N/A |
| Direct Detect® | Merck | Cat# DDHW00010-WW |
| Gradient maker | BioComp | N/A |
| Ultracentrifuge tubes | Seton tubes | Cat# S7030 |
| SW 41 Ti rotor | Beckman Coulter | Cat# 331362 |
| Piston fractionator | BioComp | N/A |
| Zentrifuge 380R | Hettich | Cat# 3220 |
| Dionex, Ultimate 3000 | Thermo Fisher Scientific | N/A |
| LUMOS mass spectrometer | Thermo Fisher Scientific | N/A |
| (LC)-MS/MS | Thermo Fisher Scientific | N/A |
| Precolumn C18 PepMap100 | Thermo Fisher Scientific | Cat# 160454 |
| C18 column | NYKKYO | Cat# NTCC-360/75-3-155) |
| Qubit Protein Assay | Invitrogen | Cat# Q33211 |
| 4 to 12% Bis-Tris Polyacrylamide gels | Invitrogen | Cat# NW04122BOX<br>Cat# WG1401BOX<br>Cat# WG1403BOX |
| Infinite M1000 Microplate Reader | Bio-Tek | N/A |
| Rotor-Gene Q instrument | QIAGEN | N/A |

## EXPERIMENTAL MODEL AND SUBJECT DETAILS

### Cell lines and culture conditions

Parental Flp-In™ 293 HEK cells (Invitrogen, Cat# R75007), standard 293 HEK cells, and all derived knock-in or knockout lines were maintained in DMEM+/+: Dulbecco’s Modified Eagle Medium (DMEM; Invitrogen, Cat# 41966052) supplemented with 10% dialyzed fetal calf serum (FCS; Bioconcept, Cat# 2-01F16-I), 25 mM HEPES (Invitrogen, Cat# 15630056), 1× MEM Non-Essential Amino Acids (Invitrogen, Cat# 11140035), and 100 U/mL penicillin–100 μg/mL streptomycin (Invitrogen, Cat# 15140122). Cells were cultured at 37°C in a humidified incubator with 5% CO₂.

Chemical compounds, including NVS1.1, cycloheximide (Focus Biomolecules, Cat# 10-117), puromycin (Sigma, Cat# P9620), and ternatin-4, were dissolved in DMSO.

## METHOD DETAILS

### Generation of knockout cell lines

USP9X knockout lines were generated using CRISPR–Cas9–mediated genome editing. A single guide RNA (sgRNA) targeting USP9X (5′- GTTGATCATGTCATCCAACT-3′) was cloned into pSpCas9(BB)-2A-Puro (PX459; GeneScript), which co-expresses SpCas9 and a puromycin resistance cassette. Flp- In™ 293 HEK cells were seeded in 6-well plates and transfected with the sgRNA-expressing vector using Lipofectamine 2000 (Invitrogen, Cat# 11668019) in Opti-MEM (Gibco, Cat# 31985062). After 24 hours, cells were selected with 2 µg/mL puromycin (Sigma, Cat# P9620) for 5 days. Surviving cells were seeded at limiting dilution into 24-well plates to isolate and raise single-cell clones. After expansion and cryopreservation, the clones were screened for gene disruption by immunoblotting and Sanger sequencing.

### Generation of knock-in cell lines

Knock-in cell lines expressing either the catalytically inactive USP9X_C1566S mutant or a silent_C1566 control were generated via CRISPR–Cas9–mediated homology-directed repair (HDR). CRISPR/Cas9 was delivered as a ribonucleoprotein (RNP) complex containing recombinant *S. pyogenes* HiFi Cas9 nuclease (125 pmol; IDT, Cat# 1081059) and Alt-R CRISPR–Cas9 sgRNA (150 pmol; IDT). The RNP complex was assembled by incubating sgRNA and Cas9 at room temperature for 20 minutes in a final volume of 5 µL. The HDR template and sgRNA were designed using the Alt-R HDR Design Tool (IDT). For electroporation, 1 × 10 cells were collected, washed once with PBS, and resuspended in 100 µL in Opti-MEM (Gibco, Cat# 31985062) per condition. The preassembled RNP (5 µL) and 250 pmol HDR template were added to the cell suspension and transferred to 2-mm electroporation cuvettes (Thermo Fisher Scientific, Cat# FB102). Electroporation was performed using the Super Electroporator NEPA21 Type II (Nepagene). Immediately after electroporation, cells were transferred to pre-warmed DMEM containing 1 µM Alt-R HDR Enhancer V2 (IDT, Cat# 10007921) and incubated for 12–24 hours at 37°C with 5% CO₂. Media were replaced with fresh DMEM+/+ and cells were cultured for an additional 24 hours. Single-cell clones were obtained by limiting dilution, expanded, and verified by Sanger sequencing.

### Generation of dTAG-USP9X knock-in cells using CRIS-PITCh v2

The dTAG-USP9X knock-in cell line was generated using the CRIS-PITCh v2 system. Genome editing was performed using a pX330-BbsI-PITCh_USP9X plasmid encoding a USP9X-specific single-guide RNA (sgRNA; 5′- TGGCTGTCATACTCGACACA-3′) together with a pCRIS-PITChv2_USP9X donor plasmid. The donor plasmid contained two 40-bp N-terminal USP9X microhomology arms flanking a puromycin resistance cassette, a T2A sequence, the FKBP12^F36V^ degron tag, and a linker region. Both plasmids were derived from pCRIS-PITChv2 and pX330-BbsI-PITCh and were kindly provided by Niels Gehring. The CRIS-PITCh v2 protocol was described previously^78,79^, and was followed here with minor modifications. Briefly, 1 × 10 Flp-In™ HEK293 cells were collected, washed once with PBS, and resuspended in 100 µL Opti-MEM (Gibco, Cat# 31985062) per condition. Of each plasmid, 1 µg was added to the cell suspension, which was then transferred to 2-mm gap electroporation cuvettes (Thermo Fisher Scientific, Cat# FB102). Electroporation was performed using a NEPA21 Type II Super Electroporator (Nepagene). Immediately after electroporation, cells were transferred to pre-warmed DMEM+/+ supplemented with 1 µM Alt-R HDR Enhancer V2 (IDT, Cat# 10007921) and incubated for 12–24 hours at 37°C with 5% CO₂. Media was then replaced with fresh DMEM+/+, and cells were cultured for an additional 24 hours. Puromycin selection (2 µg/mL; Sigma, Cat# P9620) was initiated 24 hours later and maintained for 5 days. Surviving cells were seeded at limiting dilution, and single colonies were transferred to 24-well plates 7–10 days later. Clones were expanded and screened for correct knock-in integration by PCR. Homozygous knock-in clones were subsequently treated with 500 nM dTAGV-1 (MedChemExpress, Cat# HY-145514D) to validate efficient USP9X protein degradation.

### siRNA-mediated knockdown

For siRNA transfection, ∼1 × 10 cells in 6-well plates were transfected with 50 nM siRNA targeting USP9X, EDF1, or GIGYF2, or with a non-targeting control (Lullaby reagent, Oz Biosciences, Cat# LL71000). siRNA and 14 μL of Lullaby reagent were mixed in 500 μL in Opti-MEM (Gibco, Cat# 31985062) and added to cells (final volume 2 mL per well). After 24 hours, cells were split 1:4 and transfected again under the same conditions. 24 hours after the second transfection, cells were treated with either DMSO or 25 μM NVS1.1 for 6 hours, harvested by scraping in cold PBS, pelleted at 500 × g for 5 minutes at 4°C, and stored at −80°C.

### Immunoblot

Cell pellets were lysed in RIPA buffer (50 mM Tris-HCl, pH 8.0; 150 mM NaCl; 5 mM EDTA; 1% IGEPAL CA-630; 0.5% sodium deoxycholate; 0.1% SDS). Lysates were cleared by centrifugation at 13,000 × g for 10 minutes at 4°C. Protein concentration was measured using an infrared-based system – Direct Detect^®^ (Merck, Cat# DDHW00010-WW), and 15 μg were mixed with 4× NuPAGE LDS loading buffer (Thermo Fisher Scientific, Cat# NP0008) to a final 1.5× concentration, supplemented with 25 mM DTT, and resolved on 4–12% Bis-Tris gels (Invitrogen, Cat# NW04122BOX). Proteins were transferred to nitrocellulose membranes using the Trans-Blot Turbo Transfer System (Bio-Rad, Cat# 1704150). Membranes were blocked in 5% milk in TBS containing 0.1% Tween-20 (TBS-T) and incubated with primary antibodies diluted in 5% BSA/TBS-T for 1,5 hours at room temperature or overnight at 4°C. After washing with TBS-T, membranes were incubated with IRDye-conjugated secondary antibodies for 1 hour, washed, air-dried, and scanned using the Odyssey Imaging System (LI-COR).

### Immunoprecipitation

For pulldown assays, cells (∼80% confluence in 15-cm dishes) were treated with DMSO or 25 μM NVS1.1, washed with cold PBS, and harvested by scraping. Pellets were lysed in IP buffer (50 mM HEPES, pH 7.3; 600 mM NaCl; 0.5% Triton X-100) and homogenized by centrifugation in a Zentrimix 380R (Hettich, Cat# 3220) at 1,500 rpm for 4 minutes at 5°C. Lysates were cleared by centrifugation at 13,000 × g for 10 minutes at 4°C, and total protein was quantified using Direct Detect^®^ (Merck, Cat# DDHW00010-WW) and normalized to equal protein concentrations across samples. For immunoprecipitation, lysates were incubated with 6 μg eRF1 antibody (Santa Cruz Biotechnology, Cat# sc-365686) pre-bound to 0.75 mg Dynabeads Protein G (Invitrogen, Cat# 10004D) for 1 hour at 4°C with rotation. Beads were washed three times with IP buffer, and bound proteins were eluted in 1.5× LDS sample buffer containing 25 mM DTT at 70°C for 10 minutes.

### Polysome fractionation

Cells grown in 15-cm dishes (∼80% confluence) were treated with DMSO or 25 μM NVS1.1 for 1 hour, followed by 100 μg/mL cycloheximide (Focus Biomolecules, Cat# 10-117) for 4 minutes. Cells were washed with cold PBS containing cycloheximide, scraped, and pelleted at 500 × g for 5 minutes at 4°C. Pellets were lysed in 300 μL lysis buffer (10 mM Tris-HCl, pH 7.5; 10 mM NaCl; 10 mM MgCl_2_;1% Triton X-100; 1% sodium deoxycholate; 100 μg/mL cycloheximide; 1 mM DTT; 0.1 U/μL RNase inhibitor) for 2 minutes on ice. Lysates were cleared (16,000 × g, 5 minutes, 4°C) and digested with 400 U micrococcal nuclease S7 (Thermo Fisher Scientific, Cat# EN0181) in 1 mM CaCl_2_ for 20 minutes at room temperature, followed by quenching with 1 mM EGTA. Sucrose gradients (15–50%) were prepared using a BioComp gradient maker in gradient buffer (10 mM Tris-HCl, pH 7.5; 100 mM NaCl; 10 mM MgCl_2_; 100 μg/mL cycloheximide; 1 mM DTT) and centrifuged at 40,000 rpm for 2 hours at 4°C (SW41 Ti rotor; Beckman Coulter). Gradients were fractionated using a BioComp piston fractionator, and A260 profiles were recorded. Fractions were stored at −80°C until further processing.

### Global proteome analysis by label-free mass spectrometry

For global proteome analysis following polysome fractionation, fractions corresponding to disomes or heavy polysomes were combined with one volume of acetone and 0.1 volumes of 100% trichloroacetic acid (TCA). Samples were incubated overnight at −80 °C to precipitate proteins, which were subsequently pelleted by centrifugation at 16,000 × g for 5 minutes at 4 °C. The resulting pellets were washed three times with ice-cold acetone, with centrifugation performed under the same conditions after each wash. Heavy polysome fractions were pooled, and both heavy polysome and disome samples were dried for 20 minutes in a SpeedVac concentrator before storage at −80°C.

Protein pellets were resuspended in 12 µL of lysis buffer containing 8 M urea and 100 mM Tris-HCl (pH 8). Protein concentration was determined from a 1:10 aliquot using the Qubit Protein Assay (Invitrogen, Cat# Q33211). The remaining sample was reduced and alkylated, followed by enzymatic digestion. Proteins were initially digested with LysC (Promega, Cat# VA117A) for 2 hours at 37 °C and subsequently with 100 ng of trypsin (Promega, Cat# V5111) overnight at room temperature. Peptide samples were analyzed by liquid chromatography using a Dionex Ultimate 3000 system (Thermo Fisher Scientific) coupled to a LUMOS mass spectrometer (Thermo Fisher Scientific), with two injections of 500 ng peptide per sample.

### DiGly (K-***ε***-GG) ubiquitin remnant analysis

DiGly proteomics was performed in USP9X CD and USP9X-silent control cells. Cells were grown in 15 cm dishes until reaching confluency, washed once with cold PBS, and scraped. Cell pellets were collected and lysed in lysis buffer containing 8 M urea, 150 mM NaCl, 50 mM Tris-HCl (pH 8.0), and 5 mM N-ethylmaleimide (MedChemExpress, Cat# HY-D0843), supplemented with complete protease inhibitor (TargetMol, Cat# 0001) and phosphatase inhibitor cocktails (TargetMol, Cat# C0002 and C0003). Samples were homogenized by centrifugation in a Zentrimix 380R (Hettich, Cat# 3220) at 1,500 rpm for 4 minutes at 5°C. Lysates were subsequently cleared by centrifugation at 13,000 × g for 10 minutes at 4°C, and total protein concentration was determined using Direct Detect® (Merck, Cat# DDHW00010- WW). A total of 1 mg of protein per sample was subjected to overnight digestion with trypsin (Promega, Cat# V5111) according to previously described protocols.^17^ Ubiquitin remnant-containing peptides were enriched using Ubiquitin Remnant (K-ε-GG) Immunoaffinity Beads (Cayman, Cat# 34699) according to the manufacturer’s instructions. Briefly, lyophilized peptides were dissolved in 1 mL ice-cold Immunoaffinity Purification (IAP) buffer (50 mM MOPS, pH 7.2, 10 mM disodium phosphate, 50 mM NaCl) and vortexed. The peptide supernatant was added to tubes containing immunoaffinity beads (30 µL bead volume) pre-washed four times with 1 mL of cold IAP buffer. Samples were rotated for 1 hour at 4°C, after which the supernatant was removed. Beads were washed three times with 1 mL ice-cold IAP buffer and three times with 1 mL ice-cold HPLC-grade water. Peptides were eluted by two consecutive incubations with 50 µL 0.1% trifluoroacetic acid (TFA) for 10 minutes at room temperature with gentle agitation every 2–3 minutes. Eluates were dried by vacuum concentration and stored at −80°C until analysis. Quantification of ubiquitination site occupancy was performed by summing signal intensities for modified residues per sample, normalizing to total protein intensity, and averaging across biological replicates as described previously.^18^

### Puromycin incorporation assay

Cells were seeded in 6-well plates and grown to ∼80% confluence, and were treated with 25 μM NVS1.1 for 6 hours or with DMSO or 100 μg/mL cycloheximide for 15 minutes. Cells were then incubated with 10 μg/mL puromycin (Sigma, Cat# P9620) in DMEM+/+ for 15 minutes in the continued presence of NVS1.1, DMSO, or cycloheximide. Cells were washed with ice-cold PBS and harvested by scraping. Cell pellets were collected by centrifugation (500 × g, 5 minutes, 4°C) and lysed in RIPA buffer for subsequent immunoblotting.

### Flow cytometry analysis

For the dual-fluorescence reporter assay, ∼1 × 10 cells in 6-well plates were transfected with K(AAA)₂₀, K(AAA)₀, GFP, or RFP plasmids. 48 hours post-transfection, cells were washed with PBS, trypsinized, resuspended in DMEM+/+, and centrifuged at 500 × g, 5 minutes, 4°C. Pellets were resuspended in 500 μL cold PBS, filtered through a 70 μm strainer, and analyzed using a CytoFLEX S flow cytometer (Beckman Coulter). A total of 10,000 events were collected per sample. Data were analyzed using FlowJo, with gating based on untransfected controls; GFP⁺ and RFP⁺ populations were gated separately, compensated, and quantified based on median fluorescence intensity (MFI).

### Global protein synthesis measurement via L-AHA incorporation

Global protein synthesis was quantified by L-azidohomoalanine (L-AHA) incorporation, as previously described,^80^ with minor modifications. Briefly, prior to labeling, cells (at 70–90% confluency) were washed once with PBS and incubated in methionine-free DMEM (Gibco, Cat# 21013024) supplemented with either 25 µM NVS1.1 for 4 hours or 100 µg/mL cycloheximide for 30 minutes. Following treatment, 100 µM L-AHA (Vector Laboratories, Cat# CCT-1066) was added to each well, and incubation was continued for an additional 2 hours. Cells were then washed twice with PBS and fixed with 4% paraformaldehyde (Sigma-Aldrich, Cat# 158127) for 15 minutes at room temperature, followed by permeabilization with 0.5% Triton X-100 (Sigma-Aldrich, Cat# 93443) for 15 minutes. Incorporated L-AHA was fluorescently tagged via a Click-iT reaction using 1 µM XFD488-alkyne (AAT Bioquest, Cat# 1700) in PBS containing 10 mM sodium L-ascorbate (Sigma-Aldrich, Cat# A7631) and 2 mM CuSO₄ (Sigma-Aldrich, Cat# C1297) for 30 minutes at room temperature in the dark. To minimize background fluorescence, cells were washed twice with 1% BSA (Sigma-Aldrich, Cat# 05470). Fluorescence was quantified using a CytoFLEX S flow cytometer (Beckman Coulter), collecting 10,000 events per sample. Calculation and visualization were done using FlowJo software.

### Proteasome activity assay

To assess proteasomal activity in both WT and USP9X KO cells, a proteasome assay kit (Cayman chemicals; Cat# 10008041) was utilized. The positive control, EGCG, a proteasome inhibitor, was also included in the kit. In brief, cells were seeded into 96-well microtiter plates (0,01 ×10^6^ cells per well) and allowed to adhere overnight. The plate was centrifuged at 500 × g for 5 minutes, and the culture media was removed. Assay buffer (200 µL) was added to each well, followed by another centrifugation step at 500 × g for 5 minutes, and subsequent aspiration of the supernatant. Lysis buffer (100 µL) was then added to each well, and the plates were gently shaken for 30 minutes. After centrifugation at 1,000 × g for 10 minutes, 90 µL of supernatant from each well was transferred to the corresponding wells in a black 96-well plate (Cayman). For the measurement of sample activity, assay buffer (10 µL) was added to these wells, followed by the addition of the proteasome substrate, SUC-LLVY-AMC (10 µL). Fluorescence intensity was quantified using the Tecan Infinite M1000 Microplate Reader (Bio-Tek) with excitation at 360 nm and emission at 480 nm.

### RNA analysis by RT-qPCR

Total RNA was isolated using the Maxwell RSC simplyRNA Cells Kit (Promega, Cat# AS1390) according to the manufacturer’s instructions. RNA concentration was determined by NanoDrop spectrophotometry, and 1 μg of total RNA was reverse transcribed using random hexamer primers and AffinityScript Multi-Temp Reverse Transcriptase (Agilent Technologies, Cat# 600109). Quantitative PCR reactions (15 μL) were prepared using 24 ng of cDNA, 500 nM of each primer, and 1× Brilliant III Ultra-Fast SYBR Green qPCR Master Mix (Agilent Technologies, Cat# 600883). qPCR analyses were performed on a Rotor-Gene Q instrument (QIAGEN), with all samples analyzed in technical duplicates. Control reactions lacking reverse transcriptase were included in each experiment to verify the absence of genomic

DNA contamination. Relative transcript levels were calculated using the comparative Ct (ΔΔCt) method.

### Quantification and statistical analysis

Data are presented as mean ± SD. Statistical analyses were performed using GraphPad Prism 9. Statistical significance was determined using unpaired two-tailed Student’s t-tests or one-way or two-way ANOVA followed by Tukey’s multiple-comparisons test, as indicated in the figure legends. Experiments were performed with at least three independent biological replicates, and representative data are shown.

