## Supplemental Figures S1 to S7 for "USP9X promotes the degradation of trapped translation factors on collided ribosomes"

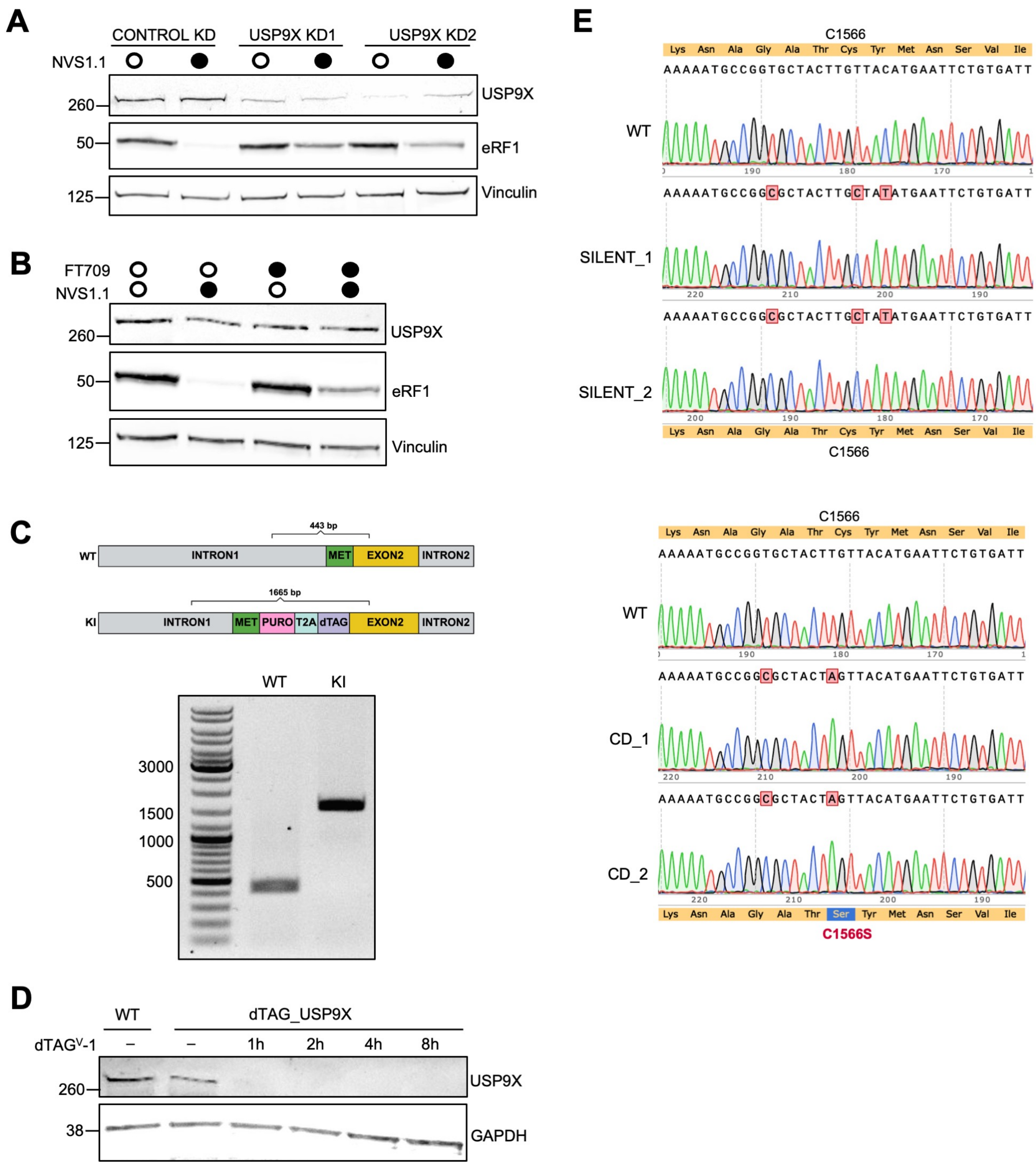


**Figure S1. USP9X depletion partially stabilizes eRF1 following NVS1.1 treatment**

**(A)** Immunoblot analysis of Flp-In™ HEK293 cells transfected with two independent USP9X-targeting siRNAs or a non-targeting control siRNA.

**(B)** Immunoblot analysis of cells treated with the USP9X inhibitor FT709. Cells were pretreated with 10 µM FT709 for 24 hours, followed by treatment with 25 µM NVS1.1 for 6 hours in the continued presence of FT709.

**(C)** Schematic representation of the *USP9X* locus in WT and KI cells. Correctly targeted clones were validated by PCR, yielding products of 443 bp for the WT allele and 1665 bp for the KI allele. PCR products were resolved on a 1% agarose gel.

**(D)** Immunoblot analysis of the positive KI clone following treatment with 500 nM dTAG$^V$-1.

**(E)** Sanger sequencing results of genomic DNA from CRISPR-Cas9 knock-in experiments generating USP9X silent C1566 and catalytic-dead USP9X C1566S cell lines. Sequences were aligned to the WT control sequence**.**

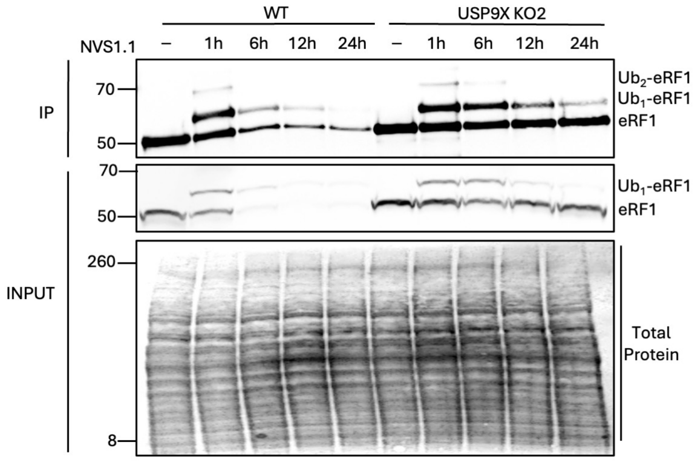


**Figure S2. Time course analysis of eRF1 ubiquitination in WT and USP9X KO cells.**

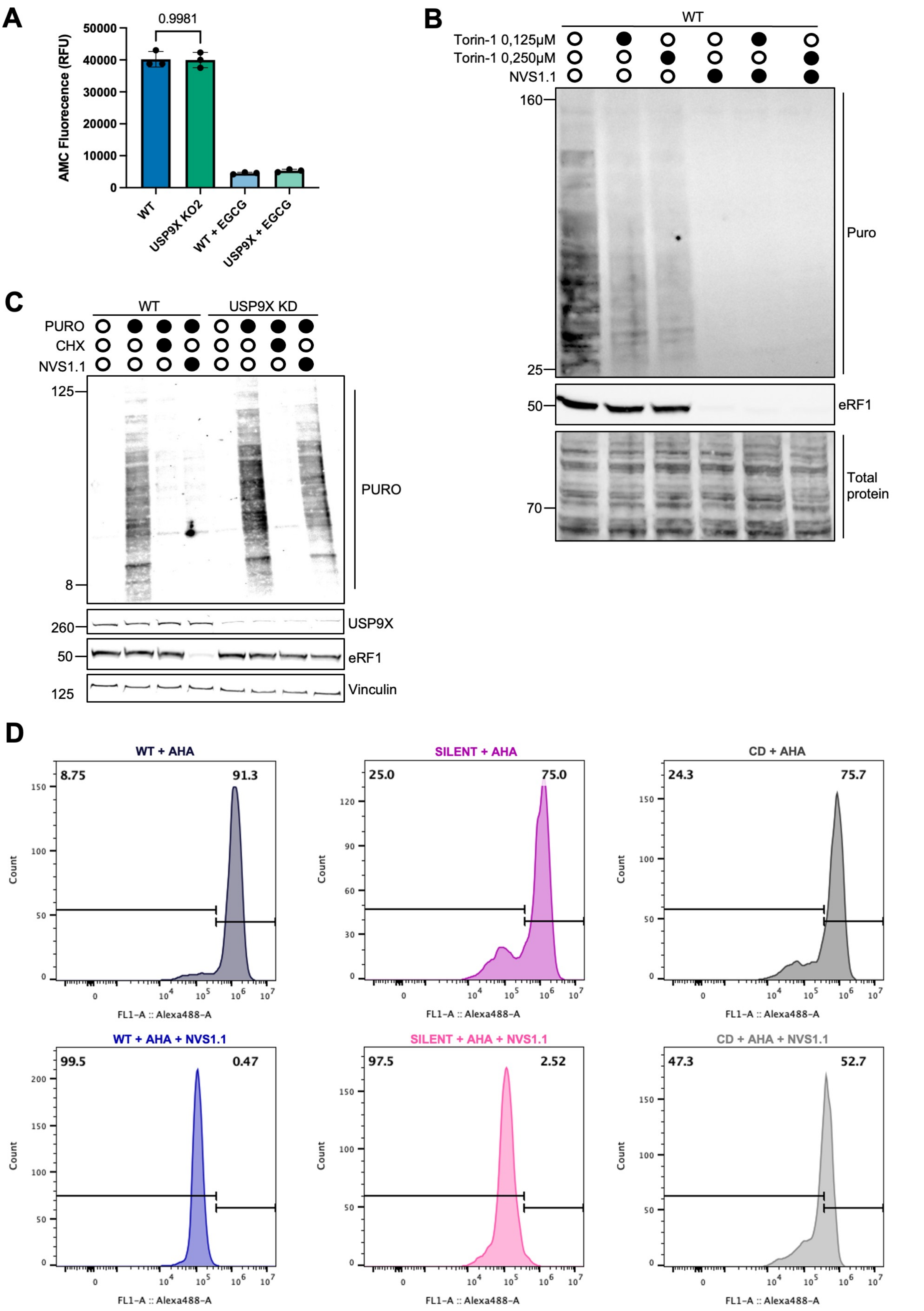


**Figure S3. USP9X depletion attenuates NVS1.1-induced translational repression**

**(A)** Fluorometric assay measuring 20S proteasome activity in WT and USP9X KO cells. Activity was assessed using the fluorogenic substrate LLVY-AMC, which releases the fluorescent product 7-amino-4-methylcoumarin (AMC) upon cleavage. AMC fluorescence was measured at an excitation wavelength of 380 nm.

**(B)** Cells were pre-treated with Torin-1 (0.125 µM or 0.25 µM) for 24 hours, followed by treatment with 25 µM NVS1.1 for 6 hours in the continued presence of Torin-1. Global translation was assessed by puromycin incorporation.
**(C)** Puromycin incorporation assay in WT and USP9X KD cells, performed as in Figure 3A.
**(D)** Quantification of L-azidohomoalanine (L-AHA) incorporation in WT, USP9X silent, and USP9X CD cells treated with 25 µM NVS1.1 or DMSO for 6 hours. L-AHA (100 µM) was added 4 hours after treatment and remained present for the final 2 hours. Bar lines indicate the gating strategy and corresponding percentages of cells without L-AHA incorporation (left) and with incorporated L-AHA (right).

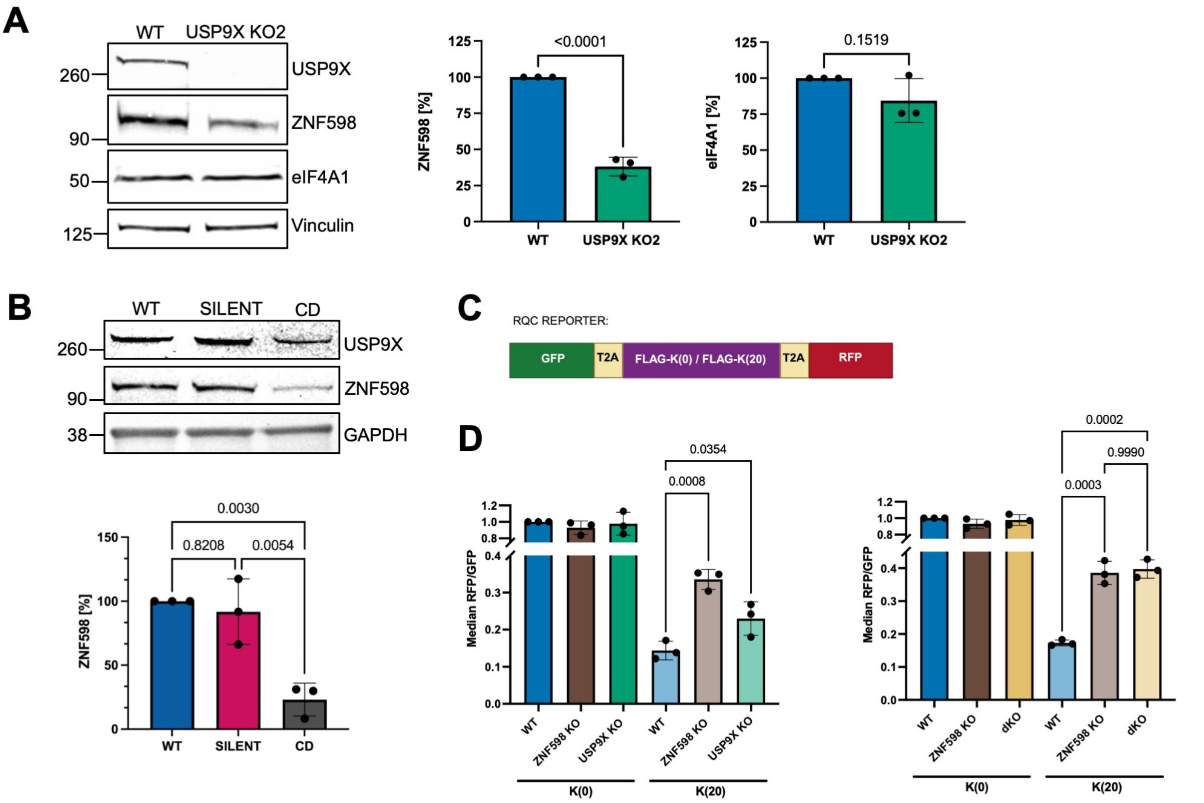


**Figure S4. USP9X loss reduces ZNF598 levels and increases poly(A)-induced readthrough**
**(A)** Immunoblot analysis and quantification of ZNF598 and eIF4A1 protein levels in WT and USP9X KO cells. Bar graphs show densitometric quantification of ZNF598 and eIF4A1 levels normalized to vinculin (mean ± SD, n = 3).
**(B)** Immunoblot analysis and quantification of ZNF598 protein levels in WT, USP9X silent, and USP9X CD cell lines. Bar graphs show densitometric quantification of ZNF598 abundance normalized to GAPDH (mean ± SD, n = 3).
**(C)** Schematic of dual-fluorescence reporter constructs. The K(20) reporter contains an internal poly(A) tract that induces ribosome stalling, whereas the K(0) reporter lacks the poly(A) sequence and serves as a control.
**(D)** Quantification of median RFP:GFP fluorescence ratios in cells expressing stalling or control reporters. Left: comparison of WT, ZNF598 KO, and USP9X KO cells. Right: comparison of WT, ZNF598 KO, and ZNF598/USP9X dKO cells. Each data point represents the median ratio from ~10,000 cells; error bars show mean ± SD (n = 3).

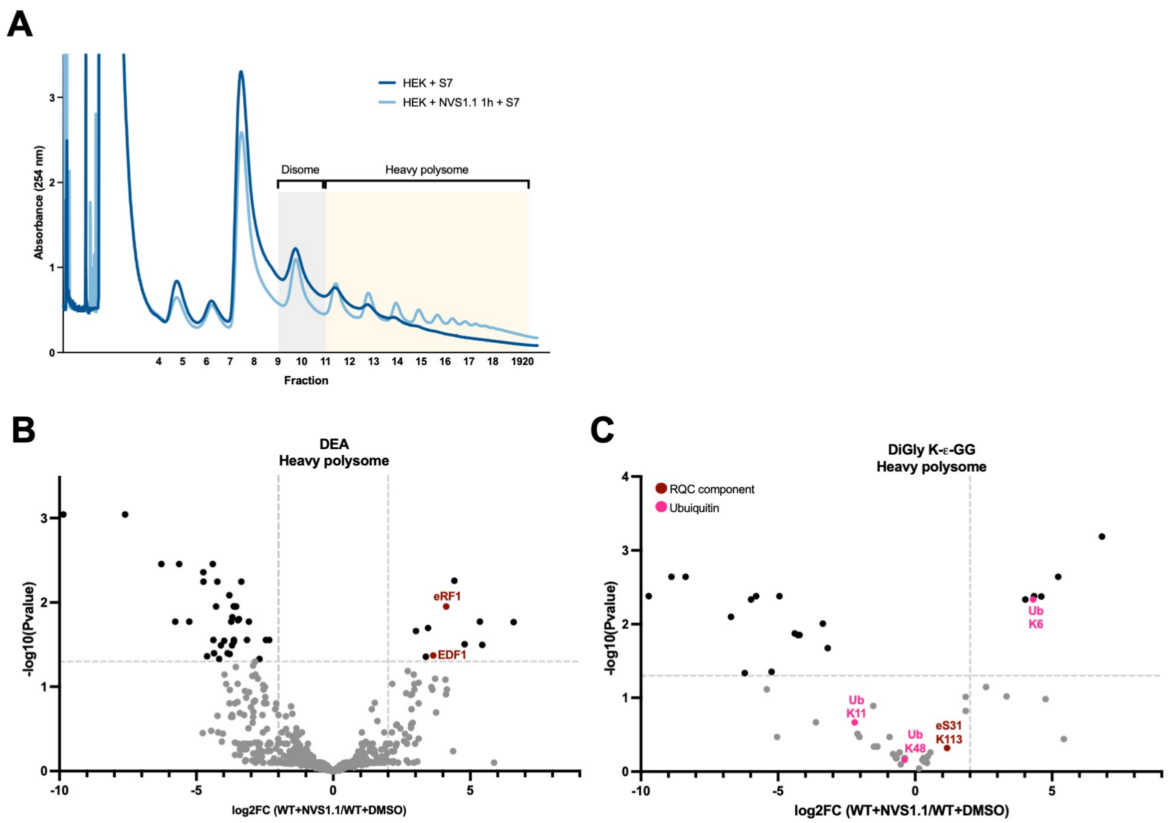


**Figure S5. eRF1 and K6-linked ubiquitination are enriched in collided ribosomes following NVS1.1 treatment.**
**(A)** Polysome profiling of cells treated with or without NVS1.1. Cells were treated with DMSO or 25 µM NVS1.1 for 1 hour, lysed, and subjected to micrococcal nuclease S7 digestion prior to separation through 15%–50% sucrose gradients. Gradients were monitored by A260, and fractions corresponding to disomes and heavy polysomes were collected for mass spectrometry.
**(B)** Differential expression analysis (DEA) identifying proteins enriched in heavy polysome fractions upon NVS1.1 treatment (from A).
**(C)** Ubiquitination analysis of proteins derived from heavy polysome fractions (from A).

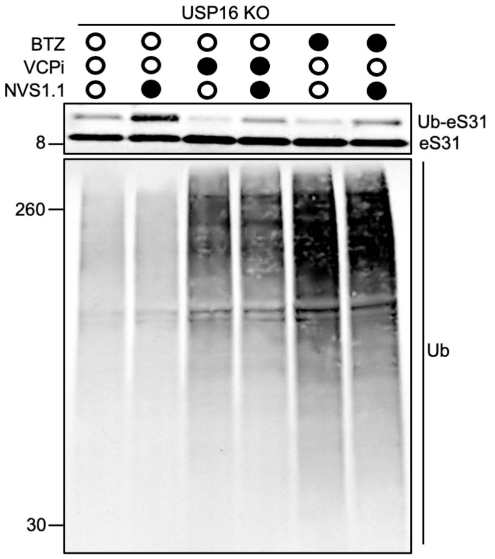


**Figure S6. VCP and proteasome inhibition do not prevent eS31 ubiquitination following NVS1.1 treatment.** USP16 KO cells were pre-treated with 10 µM VCP inhibitor (VCPi) or 0.5 µM bortezomib (BTZ) for 5 hours, followed by treatment with 25 µM NVS1.1 for 1 hour in the continued presence of the inhibitors. Immunoblot analysis of eS31 ubiquitination is shown. Total ubiquitin (Ub) accumulation confirms effective inhibition of VCP- and proteasome-dependent protein turnover.

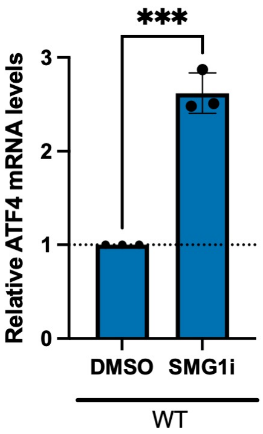


**Figure S7. NMD inhibition activates the integrated stress response.** qPCR analysis of relative *ATF4* mRNA levels in cells treated with DMSO or 0.5 μM SMG1i for 24 hours. *ATF4* expression was normalized to *ACTB*. Data represent the mean ± SD from three independent biological replicates. Statistical significance was determined using an unpaired, two-tailed Student's t-test. ***, P < 0.001.
